# No need to choose: independent regulation of cognitive stability and flexibility challenges the stability-flexibility tradeoff

**DOI:** 10.1101/2021.08.10.455850

**Authors:** Raphael Geddert, Tobias Egner

## Abstract

Adaptive behavior requires the ability to focus on a current task and protect it from distraction (*cognitive stability*) as well as the ability to rapidly switch to another task in light of changing circumstances (*cognitive flexibility*). Cognitive stability and flexibility have been conceptualized as opposite endpoints on a *stability-flexibility tradeoff* continuum, implying an obligatory reciprocity between the two: greater flexibility necessitates less stability, and vice versa. Surprisingly, rigorous empirical tests of this critical assumption are lacking. Here, we acquired simultaneous measurements of cognitive stability (congruency effects) and flexibility (switch costs) on the same stimuli within the same task, while independently varying contextual demands on these functions with block-wise manipulations of the proportion of incongruent trials and task switches, respectively. If cognitive stability and flexibility are reciprocal, increases in flexibility in response to higher switch rates should lead to commensurate decreases in stability, and increases in stability in response to more frequent incongruent trials should result in decreased flexibility. Across three experiments, using classic cued task switching (Experiments 1 and 3) and attentional set shifting (Experiment 2) protocols, we found robust evidence against an obligatory stability-flexibility tradeoff. Although we observed the expected contextual adaptation of stability and flexibility to changing demands, strategic adjustments in stability had little influence on flexibility, and vice versa. These results refute the long-held assumption of a stability-flexibility tradeoff, documenting instead that the cognitive processes mediating these functions can be regulated independently – it is possible to be both stable and flexible at the same time.

## Introduction

Adaptive behavior requires the ability to focus on a single task and protect it from distraction (*cognitive stability*), as well as the ability to rapidly switch tasks in light of changing circumstances (*cognitive flexibility*). For example, while reading a book in a busy coffeeshop, one must ignore the sounds of nearby conversations and other distractions. Conversely, reading the same book during one’s subway commute to work requires switching between focusing on the book and monitoring the stops to ensure one exits at the correct destination. As these examples illustrate, neither high stability nor flexibility is inherently preferable; rather, it is the capacity to dynamically match one’s level of stability or flexibility to optimally suit one’s current context – a marker of strategic cognitive control – that is crucial for successfully navigating different environments. Accordingly, impaired regulation of stability and flexibility has been implicated in numerous psychiatric disorders, including autism spectrum disorder (D’Cruz et al., 2013; Kenworthy et al., 2008; Poljac & Bekkering, 2012) and attention deficit hyperactive disorder (Cepeda et al., 2000; Corbett et al., 2009; Craig et al., 2016). Therefore, understanding the mechanisms mediating cognitive stability and flexibility is of vital importance to both basic and translational research.

Due to their seemingly contradictory nature, cognitive stability and flexibility have commonly been conceptualized as antagonists in an optimization problem, the so-called *stability-flexibility tradeoff* (Braem & Egner, 2018; Dreisbach & Fröber, 2019; Goschke, 2013). Inherent in this conceptualization is the assumption that stability and flexibility are opposing endpoints on a stability-flexibility continuum and are thus reciprocal – more stability necessarily comes at the cost of less flexibility, and vice versa. For example, (Goschke, 2003, 2013) proposed a single meta-control parameter (the “updating threshold”) that determines the ease with which one can switch tasks and protect a task set from distraction. When this threshold is low, task-set updating (i.e., task switching) is easy but task-set shielding is commensurately impaired. When the threshold is high, task switching is more difficult but in turn the current task set is well protected from interference. This reciprocity perspective has been a central tenet of the vast majority of literature on cognitive flexibility (Dreisbach & Fröber, 2019; Schlüter et al., 2019; Serrien & O’Regan, 2019; Siqi-Liu & Egner, 2020). However, despite its parsimony, the critical assumption of reciprocity and antagonism between stability and flexibility arguably remains unproven. The goal of the current study was to provide a stringent empirical test of the validity of this fundamental assumption.

### Evidence from Task Switching Studies

Support for the reciprocity view of cognitive stability and flexibility comes primarily from cued task switching (TS) experiments. Here, participants are usually required to switch between two or more tasks based on trial-by-trial cues (Kiesel et al., 2010; Koch et al., 2018; Vandierendonck et al., 2010). For instance, they may be cued to either classify a number as odd or even, or as greater or less than some reference value. The ubiquitous finding of a switch cost (slower and more error-prone task performance on task switches versus task repetitions) is commonly used as an indicator of one’s level of cognitive flexibility (Braem & Egner, 2018; Dreisbach & Fröber, 2019), with smaller switch costs indicating a more flexible task focus. Of particular importance to the present discussion is that switch costs often interact with a concurrent measure of cognitive stability in these experiments, namely the cross-task response congruency effect (Meiran & Kessler, 2008; Muhmenthaler & Meier, 2021). This congruency effect refers to the finding that when two tasks have to be performed with overlapping response sets, performance is slower and more error-prone for stimuli that are associated with different responses under the two task rules (incongruent stimuli) than stimuli that require the same response under both rules (congruent stimuli). Similar to switch costs, cross-task congruency effects are often interpreted to reflect a set point on the stability-flexibility continuum (Dreisbach & Fröber, 2019), with smaller congruency effects reflecting less cross-task interference, and thus a more stable task focus. Crucially, under this type of task design, the cross-task congruency effect is often found to be higher on task switch trials compared to repetition trials (Kiesel et al., 2010; Wendt & Kiesel, 2008). This interaction between task switching and cross-task congruency effects has been interpreted as supporting the reciprocity assumption of stability and flexibility, as it implies that task set shielding (cognitive stability) is weakened when people are required to switch tasks (i.e., be cognitively flexible), and vice versa.

However, a critical analysis of the switch cost – congruency interaction effect throws its support for the reciprocity view of cognitive stability/flexibility into question. First, this interaction is not ubiquitous, with some studies failing to find this effect (Li et al., 2019). Additionally, this interaction effect can be driven by floor/ceiling effects in the congruent trial task repetition condition (Gustavson et al., 2017; Marian et al., 2014; Steyvers et al., 2019) – if performance cannot get faster or more accurate in this condition, the difference between congruent task repetition and switch trials will necessarily be compressed compared to the difference between task repetition and switch trials in the incongruent conditions. Finally, and perhaps most importantly, this metric does not actually assess the reciprocity between dynamic contextual adjustments of cognitive stability and flexibility. In particular, the task switch-congruency interaction effect reflects the extent to which, given a particular level of flexibility, switch costs and congruency effects are related to one another, but it does not tap into whether strategic adjustments in cognitive stability or flexibility – the hallmark of cognitive control – are reciprocal (and thus, inter-dependent) or can occur independently.

### Evidence from Reward Studies

Other evidence for the reciprocity assumption of stability and flexibility comes from work investigating the influence of affective states and reward on cognition. Positive affect has been shown to enhance cognitive flexibility but at the cost of increased distractibility (for reviews see Goschke & Bolte, 2014 and Dreisbach & Fröber, 2019). Additionally, different types of reward appear to have dissociable influences on stability and flexibility, and cognitive control more generally. Whereas performance non-contingent rewards reduce proactive control, performance-contingent rewards have the opposite effect (Chiew & Braver, 2014; Fröber & Dreisbach, 2014, 2016b). More specifically, high performance-contingent rewards tend to promote stability in terms of proactive control (Chiew & Braver, 2014; Locke & Braver, 2008) or cue maintenance at the cost of reduced flexibility (Hefer & Dreisbach, 2017), although it has been demonstrated that changing reward prospects from one trial to the next promotes flexibility instead (Fröber & Dreisbach, 2016a; Shen & Chun, 2011), particularly when reward prospects increase. Performance-contingent rewards have also been shown to reduce switch costs (Kleinsorge & Rinkenauer, 2012). Finally, some evidence also suggests that the effect of reward on stability and flexibility is determined by the perceived effort needed to receive task rewards (Müller et al., 2007). Together, these studies are consistent with the idea of reciprocity between cognitive stability and flexibility, although they did not explicitly test for it, since they invariably examined either stability or flexibility, but not both. More stringent tests for reciprocity must necessarily involve simultaneous and independent measurements of stability and flexibility in order to ascertain whether strategic adjustments in one domain affect the other.

Such simultaneous stability and flexibility measurements have to date been only minimally employed. In one such study, (Braem, 2017) found that selectively rewarding task switches during cued task switching not only increased subsequent voluntary task switching but also increased congruency effects. This result is consistent with another cued-task switching study (Dreisbach & Wenke, 2011) demonstrating that irrelevant stimulus feature changes interacted with response repetition effects only in the case of task switches, suggesting that task shielding was reduced during switching. Similar to the switch cost – congruency interaction, however, the latter finding does not address whether strategic adjustments in cognitive stability and flexibility are reciprocal (i.e., there were no manipulations of switch rates or rewards). The former finding then is arguably the most compelling evidence in favor of reciprocity, since it demonstrates the simultaneous influence of rewarded task switching on both voluntary task switching and congruency effects. Then again, this study employed a between-subjects design and only investigated the effect of rewarding flexibility, not the effect of rewarding stability, and thus could not probe the reciprocity of stability and flexibility in both directions (Braem, 2017). A different study independently manipulated flexibility-related task parameters at a more sustained, global, versus a more transient, local level (Fröber et al., 2018). This work examined the interaction between context effects and reward prospects on stability and flexibility, and found that increasing reward prospects transiently increase flexibility even in the case of global contexts that promote sustained stability (Fröber et al., 2018). This finding has been interpreted to suggest that local and global regulation of stability and flexibility may be independent of each other, although each of these hypothesized mechanisms is nevertheless envisaged as performing a balancing act between stability and flexibility demands (Dreisbach & Fröber, 2019). In sum, work in reward and affect has in general been consistent with the reciprocity assumption of stability and flexibility, but never conclusively verified its existence, nor demonstrated a clear untethering of the two.

### Evidence from Modeling and Neuroscience

Finally, some evidence for the stability - flexibility tradeoff comes from modeling. In particular, (Musslick et al., 2018) used a recurrent neural network model to simulate the influence of a meta-control parameter (the gain of the network’s activation function) and demonstrated that various constraints on control could promote cognitive stability (i.e., decreased congruency effects) at the expense of decreased cognitive flexibility (i.e., higher switch costs), and vice versa. (Musslick et al., 2019) then analyzed this model from a dynamical system perspective, and found that by formally defining cognitive stability and flexibility in terms of attractors one could demonstrate a tradeoff between stability and flexibility. Crucially, this study also ran a behavioral experiment in human participants to examine the influence of low versus high proportion of cued task switches between groups, and found that although the switch proportion manipulation resulted in reduced switch costs in the high switch proportion group, there was no significant difference between groups in terms of congruency costs. Although there was a significant difference between groups in the gain parameter as predicted by the model, participants’ gains were significantly lower than optimal, and the manipulation of flexibility (i.e., switch proportion) did not have the expected impact on congruency effects. It thus remains relatively ambiguous whether strategic adjustments of stability and flexibility are in fact reciprocal in human behavior.

Alternatively, cognitive flexibility and stability could rely on processes that are in principle independent of each other, thus allowing for a non-reciprocal relationship. This premise implies that, in certain circumstances, one could simultaneously engage in both flexible and stable behavior. Although counterintuitive, such a situation may not be that uncommon. For example, we may need to flexibly switch between various relevant tasks (e.g., between reading a book and listening for one’s stop during a commute) while simultaneously ignoring other, unimportant distractors (e.g., nearby strangers’ conversations). In line with the idea of separable mechanisms of stability and flexibility, the neuroscience literature has pinpointed distinct loci of Dopaminergic (DA) action in mediating the protection versus updating of task sets (reviewed in Cools & D’Esposito, 2011), with DA levels in prefrontal cortex underpinning cognitive stability (Crofts et al., 2001; Durstewitz et al., 2000; Roberts et al., 1994) and DA levels in the striatum underpinning cognitive flexibility (Collins et al., 2000; Crofts et al., 2001). While DA-mediated functions in the prefrontal cortex and striatum often display an inverse pattern, a truly reciprocal relationship between the two has been considered unlikely in this literature (Cools & D’Esposito, 2011). Similarly, many clinical disorders, such as ADHD, are characterized by a combination of both inflexibility and distractibility (Barkley, 1997), suggesting that stability and flexibility rely on fundamentally distinct processes, even though they may often display an inverse relationship. Nevertheless, the above studies also fail to address directly the question of whether contextual adjustment of cognitive stability and flexibility are independent, as they typically examined these processes in isolation, considering cognitive flexibility in situations (or time points) where only shifting is required or incentivized, and stability in situations (or time points) where only shielding is necessary or incentivized (Cools et al., 2007). Moreover, this literature has also not examined the critical case of the interplay of dynamic contextual adjustments of stability and flexibility.

### The Current Study

Ultimately then, we conclude that the assumption of reciprocity, or conversely independence, between contextually appropriate levels of cognitive stability and flexibility (i.e., the stability/flexibility tradeoff) remains insufficiently tested. In the present study, we therefore sought to empirically evaluate this assumption by gauging whether contextual adaptation of cognitive flexibility reciprocally influences stability, one the one hand, and whether contextual adaptation of cognitive stability reciprocally influences flexibility, on the other hand.

To this end, we here combined classic cued task switching (Experiments 1 and 3) and attentional set shifting (Experiment 2) protocols - allowing us to simultaneously measure switch costs and congruency effects - with a well-established approach for inducing strategic adaptation of on-task stability and between-task flexibility. In particular, we independently manipulated the block-wise proportion of congruent (vs. incongruent) trials and the proportion of task switch (vs. repeat) trials. The former block-wise manipulation of congruency is known to produce the “list-wide proportion congruent” (LWPC) effect, whereby the mean congruency effect is substantially smaller when incongruent trials are frequent than when they are rare (Bugg & Chanani, 2011; Gratton et al., 1992; Logan & Zbrodoff, 1979, for reviews, see Bugg, 2017; Bugg & Crump, 2012). This observation has been interpreted as, and modeled by, a strategic up-regulation in on-task focus (i.e., cognitive stability) in response to frequent conflict from incongruent distracters (Botvinick et al., 2001; Jiang et al., 2014). Similarly, the latter, block-wise manipulation of switching is known to produce the “list-wide proportion switch” (LWPS) effect, whereby the mean switch cost is substantially reduced when switching is required more frequently (Dreisbach & Haider, 2006; Monsell & Mizon, 2006; Schneider & Logan, 2006; Siqi-Liu & Egner, 2020). Accordingly, the LWPS effect has been interpreted as reflecting strategic adjustments in switch-readiness (i.e., cognitive flexibility) to suit the changing contextual needs for more or less switching (Braem & Egner, 2018; Dreisbach & Fröber, 2019)^1^ [^1^Note that we assume these block-level effects to reflect cumulative, trial-by-trial adaptations in response to encountering more or less frequent conflict or switch requirements rather than a change in a sustained, block-level control parameter (Jiang et al., 2014; Siqi-Liu & Egner, 2020). For simplicity, we nevertheless analyze control adjustments via interaction effects between block type and trial type rather than modeling trial-by-trial learning. Importantly, for the logic of assessing the relationship between stability and flexibility, the distinction of whether changes in stability and flexibility derive from block-level versus cumulative learning effects is not critical.].

By independently varying the block-wise incidence of congruent versus incongruent, and task repeat versus switch trials, we could thus assess how these changing contextual demands, supposed to promote either stability or flexibility, affect mean switch cost and congruency effects. In line with large literatures on the LWPC and LWPS effects, we expected to observe smaller congruency effects (more stability) in blocks where incongruent trials are frequent than when they are rare, and smaller switch costs (more flexibility) in blocks where switch trials are frequent than when they are rare. Crucially, this novel design allowed us to additionally examine whether the anticipated contextual adaptation in stability (the LWPC effect) would impact switch costs (flexibility), and conversely, whether the anticipated contextual adaptation in flexibility (the LWPS effect) would impact the congruency effect (stability). If cognitive stability and flexibility were truly reciprocal, then a greater proportion of switch trials should not only decrease switch costs, due to a greater readiness to switch to the alternative task (flexibility), but should also increase cross-task congruency effects, as enhanced flexibility would be accompanied by reduced shielding of the relevant task set, and thus a greater susceptibility to interference from the stimulus features that are relevant to the alternative set. By the same token, a greater proportion of incongruent trials should not only result in smaller congruency effects, due to increased focus on the currently task-relevant stimulus features (stability), but should also increase switch costs, as enhanced focus on the relevant stimulus features would make it more difficult to release and shift that focus to the currently irrelevant stimulus features when the alternative task set is cued. By contrast, if it were possible to regulate cognitive stability and flexibility levels independently of each other, we would not expect the LWPC effect to modulate switch costs or the LWPS effect to modulate congruency effects. To preview our findings, over three experiments we observed consistent evidence against the notion of stability and flexibility being reciprocal.

## Experiment 1

As an initial test of the assumed inter-dependence between the regulation of cognitive flexibility and stability, we leveraged a classic task-switching paradigm. Here, participants are cued on each trial to apply one of two possible categorization rules to a stimulus that could be classified by either rule (i.e., a bivalent stimulus). We employed bivalent task stimuli and overlapping responses across the two task sets, such that alongside these task-switching requirements, some stimuli required the same motor response across sets (congruent stimuli) and others required the alternate response (incongruent stimuli). As in the prior literature, we treated congruency effects (incongruent minus congruent trial performance) as a measure of cognitive stability, and task switching costs (task switch minus task repeat trial performance) as a measure of cognitive flexibility. Finally, we manipulated the frequency of congruent/incongruent and repeat/switch trials over blocks of trials, allowing us to answer the critical question of whether strategically adapting stability also affects flexibility, and whether strategically adapting flexibility also affects stability.

### Methods

#### Participants

We based our target sample size on two recent task switching studies documenting robust adjustments in flexibility based on switch proportion or reward manipulations (Braem, 2017; Siqi-Liu & Egner, 2020). Sample sizes in the relevant experiments ranged from 40 to 56. We therefore targeted a sample size of 60. 65 participants completed the experiment on Amazon Mechanical Turk for financial compensation. 5 participants (7.7%) were excluded for failing to meet the accuracy criterion (>75% correct), leaving a final sample of 60 participants (43 male, 17 female; age range: 25 - 67, mean: 38.85, SD: 10.48). The study was approved by the Duke University Campus Institutional Review Board.

#### Stimuli and Procedure

Task stimuli consisted of single digits (1 – 9 excluding 5) presented centrally, surrounded by a colored rectangular frame (blue or red; **Figure 1**). Participants were required to indicate either the digit’s parity (i.e., is the digit odd or even?) or its magnitude (i.e., is the digit less or greater than 5?) based on the color of the frame that surrounded the number. Color-to-task mapping was randomized across participants. Participants used their left and right index fingers placed on the ‘Z’ and ‘M’ keys, respectively, to respond to both tasks. Category-response mapping (i.e., ‘Z’ = odd, ‘M’ = even and vice versa; ‘Z’ = less than 5, ‘M’ = greater than 5 and vice versa) was randomized across participants. Each trial began with a fixation cross (500 ms), followed by stimulus presentation. Stimulus presentation lasted for 1500 ms or until a response was recorded. Each trial was followed by an intertrial screen for 1200 – 1400 ms (jittered in 50 ms increments) that provided participants with feedback on their response (“Correct”, “Incorrect” or “Too Slow” if they failed to respond within 1500 ms).

**Figure 1:**
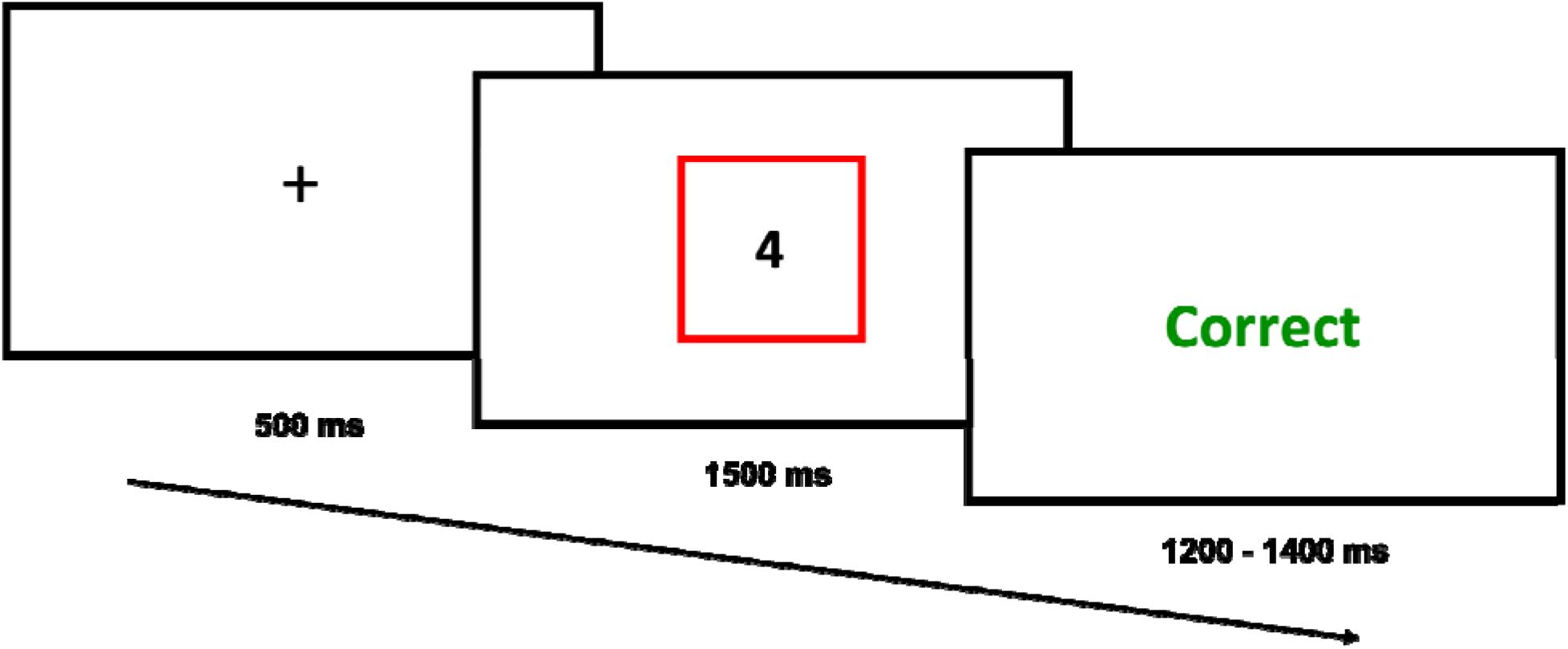
The trial structure for Experiment 1 consisted of an intertrial fixation cross preceding stimulus presentation (1-9 excluding 5), with a colored rectangle (blue or red) cueing participants to perform either the magnitude or parity task, followed by accuracy feedback.

Because the response keys for the magnitude and parity tasks overlapped, each trial could be classified as either congruent or incongruent, based on whether the keys for both tasks for a given stimulus matched or not. For example, if the parity task mapped even digits to the ‘Z’ key and odd digits to the ‘M’ key, and if the magnitude task mapped low digits to the ‘Z’ key and high digits to the ‘M’ key, then low even digits (i.e., 2 or 4) would be congruent whereas high even digits (i.e., 6 or 8) would be incongruent. The cross-task congruency effect (slower and more error-prone performance on incongruent than congruent trials) served as a measure of cognitive stability, with smaller congruency effects indicating a more stable, better shielded, task focus. Trials were also classified as repeat or switch trials based on whether or not the task cue changed between trials (i.e., a parity trial following a magnitude trial would be a switch trial). The switch cost (slower and more error-prone performance on switch than repeat trials) served as a measure of cognitive flexibility, with smaller switch costs indicating a greater readiness to switch, or a more flexible state.

The experiment began with a practice phase consisting of 3 blocks of 16 trials each. The first practice block consisted entirely of parity trials and the second of magnitude trials, with the order of those two practice blocks being randomized over participants. The third practice block consisted of a mix of parity and magnitude task trials, as in the main task. Participants were required to repeat each practice block until they achieved at least 75% accuracy. The main experiment consisted of 4 blocks of 128 trials each. Importantly, we varied the relative frequency of incongruent (vs. congruent) stimuli and the frequency of switch (vs. repeat) trials across blocks. Specifically, blocks could consist of either 25% or 75% incongruent stimuli combined with either 25% or 75% switch trials, resulting in 4 possible switch/congruency proportion combinations. The block order was counterbalanced across participants using a Latin square design. These frequency manipulations allowed us to examine strategic adaptation in performance to independently changing stability and flexibility demands via the LWPC and LWPS effects.

#### Design and Analysis

The experiment corresponded to a 2 (task sequence: switch vs. repeat) x 2 (stimulus congruency: congruent vs. incongruent) x 2 (switch proportion: 25% vs. 75%) x 2 (incongruency proportion: 25% vs. 75%) factorial design. With 128 trials per block, this meant that there were 8 trials in the rarest condition (e.g., congruent, repeat trials in the 75% incongruent, 75% switch condition) and 72 in the most common condition (e.g., incongruent, switch trials in the 75% incongruent, 75% switch condition). Importantly, the analyses that evaluated the effects of interest to our hypotheses – the two-way interactions between the list-wide proportion congruency/switch factors and trial types – involved cell sizes of 192, 192, 64, and 64 trials. The main dependent variable of interest was response time (RT) as an index of cognitive processing efficiency. We therefore ran an omnibus repeated measures analysis of variance (rmANOVA) of subjects’ mean RT with the independent variables of task sequence (switch vs. repeat), stimulus congruency (congruent vs. incongruent), block-wise switch proportion (25% vs. 75%), and block-wise incongruency proportion (25% vs. 75%). We expected to observe standard congruency effects (a main effect of stimulus congruency) and switch costs (a main effect of task sequence), as well as a LWPC effect (an interaction between stimulus congruency and congruency proportion) and a LWPS effect (an interaction between task sequence and switch proportion). In order to test our primary question regarding the interdependency of stability and flexibility, we tested whether the effect of stimulus congruency was modulated by switch proportion and whether the effect of task sequence was modulated by congruency proportion. Although only of secondary interest, we also examined higher order three-way interactions that might reveal additional nuances in the relationship between stability and flexibility.

Additionally, in order to test the strength of theoretically meaningful null effects (notably, the possible absence of the interactions predicted by a stability-flexibility tradeoff), we also ran an identical Bayesian repeated measures ANOVA in JASP (Version 0.15, JASP Team, 2021). Bayesian techniques offer numerous advantages over frequentist approaches, including the ability to interpret evidence in favor of null hypotheses (Dienes, 2014) and are now commonly considered superior to traditional frequentist approaches (Rouder et al., 2009, 2012; Wagenmakers, 2007). We used the default JASP prior for fixed effects, which is a weakly-informative prior described by a Cauchy distribution centered around zero and with a width parameter of 0.5 (Rouder et al., 2012). This corresponds to a probability of 70% that the effect size lies between -1 and 1, which is considerably more conservative than the implicit uniform prior used in conventional frequentist analyses, as the latter places equal weight on both small and extremely large effect sizes. Inverse bayes factors (BF_01_) were computed for each candidate model relative to the best-fitting model and indicate the likelihood ratio of marginal likelihoods (Kass & Raftery, 1995). Performance accuracy was only of secondary interest but was also fully analyzed, identically to RT.

#### Transparency and Openness

We report how we determined our sample size, all data exclusions, all manipulations and experimental measures, and follow JARS (Kazak, 2018). All data and task/analysis code are available at https://github.com/rmgeddert/stability-flexibility-tradeoff. Data were analyzed using R, version 4.0.2 (R Core Team, 2020) and JASP (JASP Team, 2021). The study’s design and analysis were not preregistered.

### Results

#### Trial exclusions

For the accuracy analysis, we analyzed all trials excluding practice trials and the first trial of each block (since these can be neither switch nor repeat trials). For the reaction time (RT) analysis, we further excluded all incorrect trials (7.9% of trials) and any trials where participants responded greater than 3 times the standard deviation from the mean of their response RTs (0.6% of remaining trials). We also excluded all trials where participants responded faster than 300 ms and slower than 1500 ms (<0.1% of remaining trials).

#### RT Analysis

The mean RT for each cell of the experimental design is displayed in **Figure 2a**, and descriptive statistics can be found in **Table S1**. We observed a significant main effect of task sequence (i.e., switch cost), as reflected in slower RTs for switch trials (M_switch_ = 879 ms, 95% CI [850, 908]) compared to repeat trials (M_repeat_ = 797 ms, 95% CI [768, 826]; F(1, 59) = 149.72*, p* < .001, *η_p_*^2^ = .717), and a significant main effect of stimulus congruency, due to slower RTs for incongruent (M_incongruent_ = 872 ms, 95% CI [843, 901]) compared to congruent stimuli (M_congruent_ = 804 ms, 95% CI [775, 833]; F(1, 59) = 98.91, *p* < .001, *η_p_*^2^ = .626). There was also a significant interaction between task sequence and stimulus congruency (F(1,59) = 7.82, *p* = .007, *η_p_*^2^ = .117), as congruency effects were larger on switch (M_congruencyeffect_ = 77 ms, 95% CI [62, 92]) compared to repeat trials (M_congruencyeffect_ = 60 ms, 95% CI [45, 74]), as occasionally reported in the literature (e.g., Kiesel et al., 2010; Wendt & Kiesel, 2008). Finally, we also observed main effects of both LWPS (F(1,59) = 22.07, *p* = .001, *η_p_*^2^ = .272) and LWPC (F(1,59) = 5.39, *p* = .024, *η_p_*^2^ = .084), due to slower RTs in the 75% switch condition (M_75_ = 851, 95% CI [822, 880]) than the 25% switch condition (M_25_ = 825, 95% CI [797, 854]) and in the 75% incongruency condition (M_75_ = 848, 95% CI [818, 877]) than the 25% incongruency condition (M_25_ = 828, 95% CI [799, 858]), respectively.

**Figure 2:**
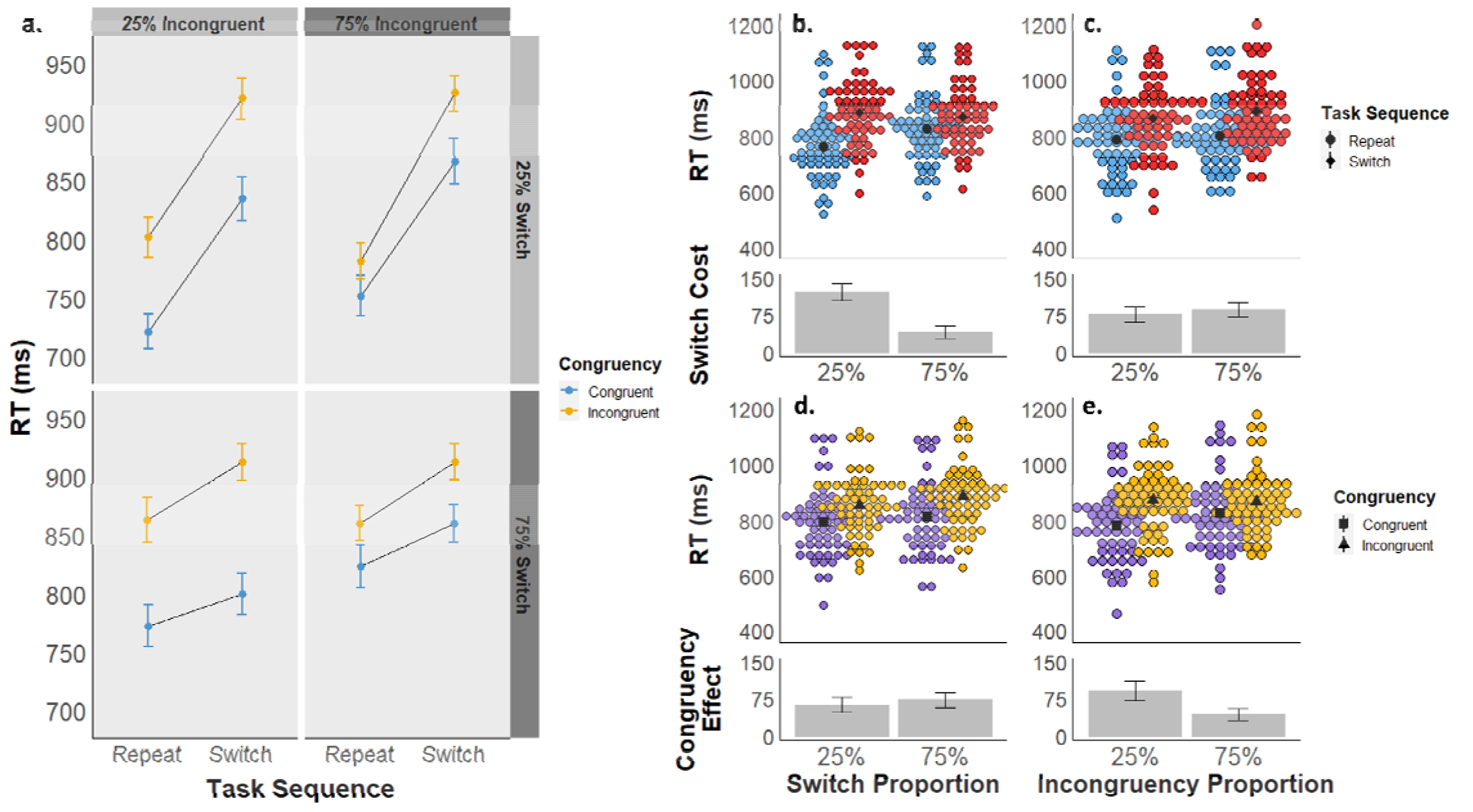
**a)** Experiment 1 mean reaction times are displayed as a function of task sequence (switch vs. repeat), stimulus congruency (congruent vs. incongruent), switch proportion (25% vs. 75%), and incongruency proportion (25% vs. 75%). **b-e)** Experiment 1 mean RTs are displayed as a function of task sequence (panels b and c) and congruency (d and e), collapsed across block-wise switch proportions (b and d) and block-wise congruency proportions (c and e). The upper graphs in each panel depict mean RTs by subject for each condition, and the lower graphs depict the mean RT difference between conditions.

Importantly, we also observed evidence for strategic adaptation in switch-readiness and task-focus. First, we found a significant interaction between task sequence and the block-wise switch proportion (i.e., 25% vs. 75% switch trials; F(1,59) = 116.09, *p* < .001, *η_p_*^2^ = .663), i.e., the LWPS effect, as switch costs were smaller in the 75% switch proportion condition (M_switchcost_= 41 ms, 95% CI [26, 56]) than in the 25% switch proportion condition (M_switchcost_ = 122 ms, 95% CI [107, 138]; **Figure 2b**). Second, we also observed a significant interaction between congruency and the block-wise congruency proportion (F(1,59) = 27.38, *p* < .001, *η_p_*^2^ = .317), i.e., the LWPC effect, as congruency effects were smaller in the 75% incongruency proportion condition (M_congruencyeffect_ = 44 ms, 95% CI [28, 60]) than in the 25% incongruency proportion condition (M_congruencyeffect_ = 92 ms, 95% CI [76, 109]; **Figure 2e**). These results replicate a large body of prior findings in the literature, suggesting that adaptive cognitive control processes enable increased cognitive flexibility (i.e., decreased switch costs) in contexts of frequent switching and increased cognitive stability (i.e., decreased distractibility) in contexts case of frequent incongruent trials.

Our primary effect of interest was the interaction between block-wise switch proportion and congruency effects, and conversely, between block-wise congruency proportion and switch costs. We found neither of these interactions to be significant. There was no interaction between the switch proportion and congruency (F(1,59) = 2.02, *p* = .161, *η_p_*^2^ = .033; **Figure 2d**). A Bayesian repeated measures ANOVA found the best-fitting model to be *task_sequence + congruency + LWPS + LWPC + task_sequence * LWPS + congruency * LWPC*. Bayesian model comparison of the congruency * LWPS interaction found a BF_01_ value of 7.1, indicating that there was more than 7 times more evidence for the model without this interaction effect, indicating medium to strong evidence in support of the null hypothesis. Likewise, there was also no significant interaction between congruency proportion and task sequence (F(1,59) = 1.74, *p* = .192, *η_p_*^2^ = .029), with a BF_01_ = 6.7 relative to the best-fitting model (above) without this interaction term (**Figure 2c**). Together, these analyses show robust evidence in support of the null hypothesis of no interaction between the LWPC factor and switch costs nor between the LWPS factor and the congruency effect. Finally, none of the other interactions in the 4-way rmANOVA were significant either (all *p* ≥ .146).

#### Accuracy Analysis

Mean accuracy for each condition is displayed in **Figure S1** and listed in **Table S2**. Accuracy was subject to ceiling effects, such that the distribution of participants’ scores violated assumptions of normality (Shapiro-Wilk W = 0.81, *p* < .001) and homogeneity (Levene’s Test F = 18.8, *p* < .001) of variance. Results should therefore be interpreted with caution. There was a significant main effect of task sequence, with participants responding more accurately on repeat (M_repeat_ = 92.5%, 95% CI [91.2, 93.7]) compared to switch trials (M_switch_ = 88.9%, 95% CI [87.3, 90.6]; F(1, 59) = 30.35, *p* < .001, *η_p_*^2^ = .340), as well as a significant main effect of stimulus congruency, with participants responding more accurately on congruent (M_congruent_ = 94.8%, 95% CI [93.8, 95.8]) compared to incongruent trials (M_incongruent_ = 86.7%, 95% CI [84.7, 88.6]; F(1,59) = 81.15, *p* < .001, *η_p_*^2^ = .579). The interaction between task sequence and congruency was significant as well (F(1,59) = 5.48, *p* = .023, *η_p_*^2^ = .085), as congruency effects were larger on switch (M_congruencyeffect_ = 9.3%, 95% CI [7.2, 11.3]) compared to repeat trials (M_congruencyeffect_ = 7.0%, 95% CI [4.9, 9.0]). There was also a main effect of switch proportion (F(1,59) = 4.87, *p* = .031, *η_p_*^2^ = .076) due to higher accuracy in the 25% switch condition (M_25_ = 91.3%, 95% CI [89.9, 92.8]) than the 75% switch condition (M_75_ = 90.1%, 95% CI [88.7, 91.5]), but the main effect of congruency effect was not significant (F(1,59) = 3.17, *p* = .080, *η_p_*^2^ = .051).

As in the RT analysis, there were significant interactions between block-wise switch proportion and task sequence (F(1,59) = 19.25, *p* < .001, *η_p_*^2^ = .246) and between block-wise congruency proportion and stimulus congruency (F(1,59) = 38.86, *p* < .001, *η_p_*^2^ = .397). Specifically, accuracy switch costs were smaller for the 75% switch condition (M_switchcost_ = 1.6%, 95% CI [0.02, 3.1]) compared to the 25% switch condition (M_switchcost_ = 5.4%, 95% CI [3.9, 6.9]), and the congruency effect was smaller for the 75% incongruent condition (M_congruencyeffect_ = 4.3%, 95% CI [2.2, 6.5]) compared to the 25% incongruency condition (M_congruencyeffect_ = 11.9%, 95% CI [9.8, 14.1]). Unlike the RT analysis, there was a significant interaction between the block-wise switch proportion and stimulus congruency, with the congruency effect being larger for the 75% switch proportion (M_congruencyeffect_ = 9.5%, 95% CI [7.5, 11.5]) compared to the 25% switch proportion condition (M_congruencyeffect_ = 6.8%, 95% CI [4.7, 8.8]; F(1,59) = 8.39, *p* = .005, *η_p_*^2^ = .125). A Bayesian rmANOVA found a best-fitting model of *task_sequence + congruency + LWPC + LWPS + task_sequence * LWPS + congruency * LWPC + congruency * LWPS*, also containing this significant interaction. Compared to this best-fitting model, the model without this term (as in the RT analysis) had a BF_01_ = 1.5, suggesting only anecdotal evidence in favor of the model with this interaction term. The interaction between task sequence and congruency proportion was not significant (F(1,59) = 0.14, *p* = .709, *η_p_*^2^ = .002), with the best model with this term having a BF_01_ = 8.2, suggesting strong evidence against this interaction. None of the other interactions were significant (all *p* ≥ .166).

### Discussion

Our primary effects of interest for Experiment 1 were the interaction between block-wise switch proportion and congruency effects, and conversely, between block-wise congruency proportion and switch costs in response time data. If the classically held assumption of a cognitive stability/flexibility tradeoff were true, increased cognitive flexibility induced by frequent switching should also increase the distracting impact of incongruent stimuli, and likewise, increased cognitive stability induced by frequent incongruent stimuli should increase switch costs. Critically, we found little evidence for inter-dependence between adaptations in stability and flexibility. Although there were numerical differences in switch costs between congruency proportions and in congruency effects between switch proportions in the direction predicted by a stability-flexibility tradeoff, these differences were not reliable as indicated by the conventional ANOVA, and the corresponding interaction terms were refuted by Bayesian analyses. Note that this was not due to a lack of power for detecting two-way interaction effects, as we found substantial evidence for interactions between congruency proportions and congruency effects (the LWPC effect) and task sequence proportions and switch costs (the LWPS effect). These results indicate that increases in task focus towards a particular task set (protecting it from cross-task interference) has no effect on one’s ability to nevertheless switch tasks, and likewise, that being more ready to switch tasks has no effect on how much other tasks interfere with the currently active task set. The accuracy analysis recapitulated the above effects but also included an interaction between task sequence and congruency proportion; however, the Bayesian rmANOVA was unable to differentiate between models with this term and without. Thus, this interaction should be interpreted cautiously, especially given that accuracy data also violated key distributional assumptions of the ANOVA. Overall, the results of Experiment 1 thus suggest that the regulation of cognitive stability and flexibility do not seem to be inversely yoked to each other.

## Experiment 2

Experiment 1 assessed the inter-dependence of regulating stability and flexibility in a classic task switching protocol, where cognitive flexibility is operationalized as switching between different task rules. In Experiment 2, we sought to investigate whether the independence between control over stability and flexibility that we observed in the context of task switching would replicate in (and thus, generalize to) another common probe of cognitive flexibility, namely, attentional set shifting (Dias et al., 1996; Robbins, 2007). In set shifting experiments, participants are cued to shift attention between different features of a compound stimulus to guide their response, but unlike in task switching, the basic task rule remains the same. One example is the classic Navon (or global-local) task (Navon, 1977, 2003), where participants are presented with large (“global”) letter shapes made up of small (“local”) letters, which can be either congruent or incongruent with the global shape (**Figure 3**). On each trial, participants are cued to classify the stimulus based on either the global or local feature. Thus, trials where the relevant feature differs from the previous trial require a shift in attentional set, either from focusing on global to focusing on local information, or vice versa, and this shift cost represents an index of cognitive flexibility. Moreover, given the bivalent nature of the stimuli, combined with overlapping response sets, the irrelevant stimulus feature in this task can produce strong interference (congruency) effects, which can in turn be used as an index of cognitive stability. In Experiment 2, we therefore adopted the Navon task as a means to manipulate demands on, and measure effects of, cognitive stability and flexibility in the domain of attentional set shifting.

**Figure 3:**
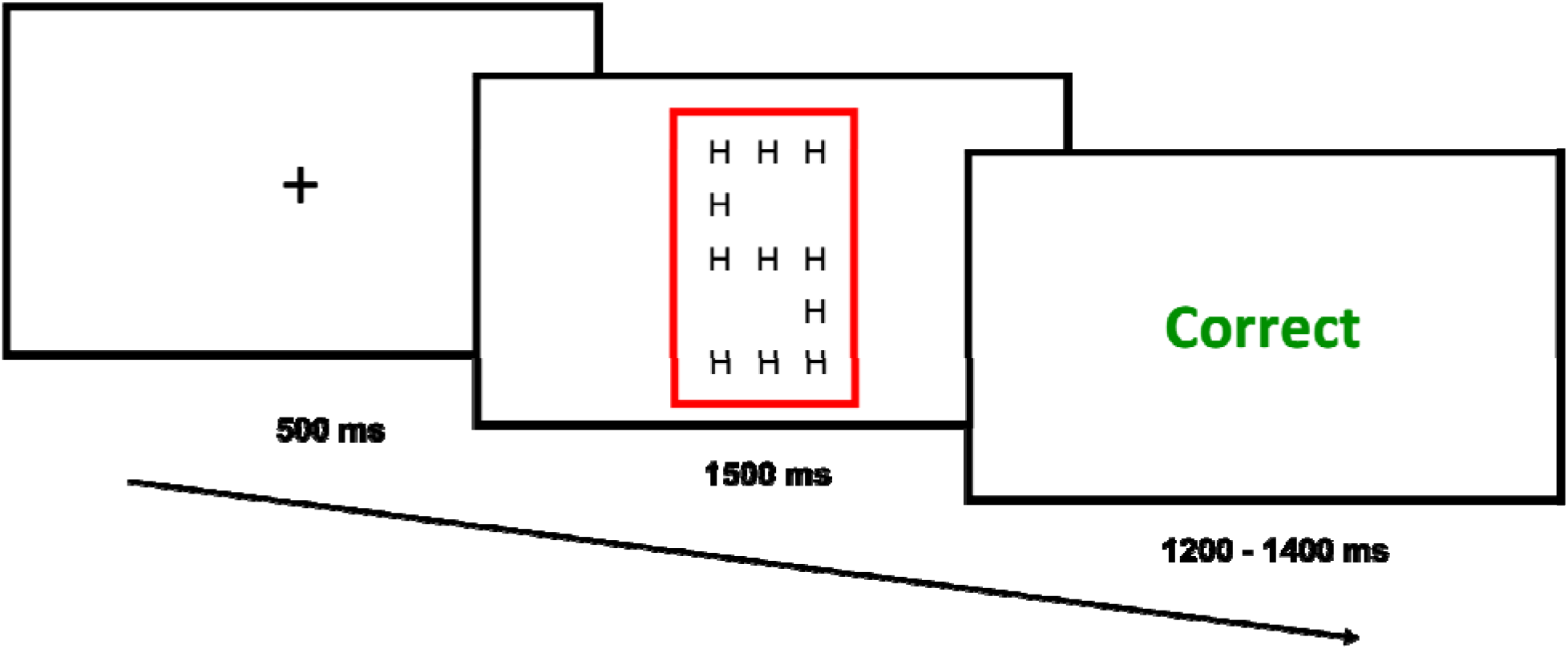
The trial structure for Experiment 2 consisted of an intertrial fixation cross preceding stimulus presentation (a large ‘S’ or ‘H’, made of smaller ‘S’s or ‘H’s), with a colored rectangle (blue or red) cueing participants to identify the global or local letter, followed by accuracy feedback.

### Methods

#### Participants

70 participants completed the experiment on Amazon Mechanical Turk for financial compensation. 9 participants (12.9%) were excluded for failing to meet the accuracy criterion (>75% correct), as well as one participant who did not have any correct trials in one of the rmANOVA cells for the RT analysis, leaving a final sample of 60 participants (male = 29, female = 31; age range: 25 - 69, mean: 38.55, SD: 11.65). The study was approved by the Duke University Campus Institutional Review Board.

#### Stimuli and Procedure

Task stimuli for Experiment 2 consisted of “Navon” stimuli: large letters, either ‘S’ or ‘H’, made of smaller letters (also ‘S’s or ‘H’s) (**Figure 3**), and participants were required to classify either the large shape or the small letters as ‘S’ or ‘H’. Participants were cued whether to respond to the large or small letters by a colored rectangle (blue or red) that surrounded each stimulus. Color-to-task mapping was randomized across participants. Participants used their left and right index fingers placed on the ‘1’ and ‘0’ keys, respectively, to indicate whether the large or small letters were an ‘S’ or ‘H’. Letter-response mapping (i.e., ‘1’ = ‘S’, ‘0’ = ‘H’, and vice versa) was randomized across participants. Each trial began with a fixation screen (500 ms), followed by task stimulus presentation (1500 ms or until a response was recorded). Each task stimulus was followed by an intertrial screen for 1200 – 1400 ms (jittered in 50 ms increments) that provided participants with feedback on their response (“Correct”, “Incorrect” or “Too Slow” if they failed to respond within 1500 ms).

Each trial was classified as being either congruent or incongruent based on whether the small and large letters matched, and as a repeat or switch trial based on whether the task cue changed between trials (i.e., a ‘large shape’ trial following a ‘small letters’ trial would be a switch trial). The congruency effect again served as a measure of cognitive stability, with smaller congruency effects indicating a more stable task focus. Likewise, the switch cost again served as a measure of cognitive flexibility, with smaller switch costs indicating a more flexible state. The practice phase consisted of 3 blocks of 16 trials each. In the first two practice blocks, participants practiced identifying first the identity of the large shape only and then the identity of the small letters only (order randomized across participants). In the third practice block, participants had to use the colored rectangular frame to determine whether to categorize the large or small letters as ‘S’ or ‘H’. The main experiment consisted of 4 blocks of 128 trials each, resulting in the same trial counts as Experiment 1. We again varied the relative frequency of incongruent stimuli (25% vs 75%) and the frequency of switch trials (25% vs 75%) across blocks, with block order counterbalanced across participants using a Latin square design.

The design and analysis strategy for Experiment 2 were identical to those of Experiment 1. We expected to again reproduce standard congruency, proportion congruent, and task switch and proportion switch effects. The main question of interest was whether congruency effects would interact with the proportion switch factor and whether switch costs would interact with the proportion congruent factor.

### Results

#### Trial Exclusions

Trial exclusion criteria for Experiment 2 were the same as for Experiment 1, including removal of incorrect trials (10%), 3 times standard deviation RT trials (0.5%), and trials faster than 300 ms or slower than 1500 ms (<0.1%).

#### RT Analysis

The mean RT for each cell of the experimental design is displayed in **Figure 4a**, and descriptive statistics can be found in **Table S1**. As expected, there was a significant main effect of task sequence, as seen in slower RTs for switch trials (M_switch_ = 899 ms, 95% CI [868, 930]) compared to repeat trials (M_repeat_ = 812 ms, 95% CI [781, 843]; F(1, 59) = 225.18, *p* < .001, *η_p_*^2^ = .792), and a significant main effect of stimulus congruency, as seen in slower RTs for incongruent (M_incongruent_ = 909 ms, 95% CI [878, 940]) compared to congruent stimuli (M_congruent_ = 802 ms, 95% CI [772, 833]; F(1, 59) = 266.26, *p* < .001, *η_p_*^2^ = .819). The interaction between task sequence and stimulus congruency was not significant (F(1,59) = 2.26, *p* = .138, *η_p_*^2^ = .037), unlike Experiment 1. There were also significant main effects of LWPS (F(1,59) = 34.19, *p* < .001, *η_p_*^2^ = .367) and LWPC (F(1,59) = 20.82, *p* < .001, *η_p_*^2^ = .261), due to slower RTs in the 75% switch condition (M_75_ = 871, 95% CI [840, 902]) than the 25% switch condition (M_25_ = 840, 95% CI [809, 871]) and in the 75% incongruency condition (M_75_ = 874, 95% CI [843, 906]) than the 25% incongruency condition (M_25_ = 837, 95% CI [806, 868]), respectively.

**Figure 4:**
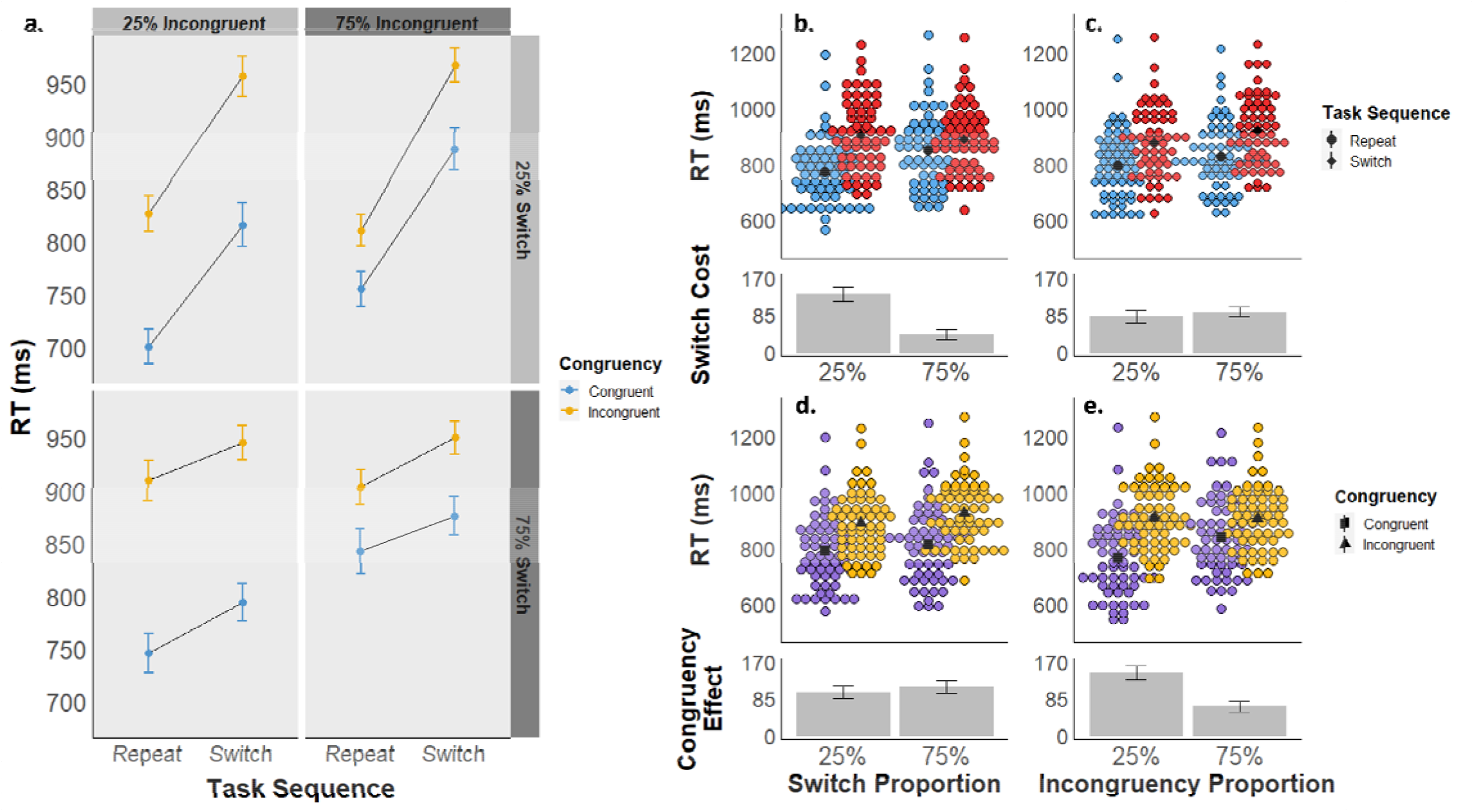
**a)** Experiment 2 mean reaction times are displayed as a function of task sequence (switch vs. repeat), stimulus congruency (congruent vs. incongruent), switch proportion (25% vs. 75%), and incongruency proportion (25% vs. 75%). **b-e)** Experiment 2 mean RTs are displayed as a function of task sequence (panels b and c) and congruency (d and e), collapsed across block-wise switch proportions (b and d) and block-wise congruency proportions (c and e). The upper graphs in each panel depict mean RTs by subject for each condition, and the lower graphs depict the mean RT difference between conditions.

We also replicated the standard LWPS and LWPC effects. Specifically, we again found a significant interaction between task sequence and the block-wise switch proportion (F(1,59) = 102.29, *p* < .001, *η_p_*^2^ = .634). Switch costs were smaller in the 75% switch proportion condition (M_switchcost_= 41 ms, 95% CI [26, 56]) than in the 25% switch proportion condition (M_switchcost_ = 134 ms, 95% CI [119, 148]; **Figure 4b**). We also observed a significant interaction between congruency and the block-wise congruency proportion (F(1,59) = 104.83, *p* < .001, *η_p_*^2^ = .640). Congruency effects were smaller in the 75% incongruency proportion condition (M_congruencyeffect_ = 67 ms, 95% CI [52, 82]) than in the 25% incongruency proportion condition (M_congruencyeffect_ = 145 ms, 95% CI [130, 160]; **Figure 4e**).

Our primary effects of interest were again the interaction between block-wise switch proportion and congruency effects, and conversely, between block-wise congruency proportion and switch costs. We found neither of these interactions to be significant. First, there was no interaction between congruency and switch proportion (F(1,59) = 3.69, *p* = .060, *η_p_*^2^ = .059; **Figure 4d**). A bayesian rmANOVA found a best fitting model of *task_sequence + congruency + LWPS + LWPC + task_sequence * LWPS + congruency * LWPC*, replicating the best fitting model from Experiment 1. Compared to this model, a model with the added congruency * LWPS interaction had a BF_01_ = 5.2, suggesting moderate evidence in favor of the null model. Likewise, there was also no significant task sequence * LWPC interaction (F(1,59) = 3.16, *p* = .081, *η_p_*^2^ = .051; **Figure 4c**), with a BF_01_ = 6.4. Together, these analyses again show robust evidence in support of the null hypothesis of no interaction between the congruency proportion and switch costs or between switch proportion and the congruency effect.

Unlike Experiment 1, the rmANOVA also detected a three-way interaction of congruency * LWPS * LWPC (F(1,59) = 5.63, *p* = .021, *η_p_*^2^ = .087), due to a smaller effect of congruency proportion on congruency effects in the 25% switch condition (M = 66 ms, 95% CI [48, 84]) than in the 75% switch condition (M = 90 ms, 95% CI [72, 108]). Note that this interaction runs opposite to what one would expect to observe based on a stability-flexibility tradeoff, with smaller LWPC effects on congruency (lower stability) in the 25% switch condition (low flexibility). Regardless, the best fitting Bayesian rmANOVA models with this significant three-way interaction term had BF_01_ = 65.8, suggesting very strong evidence in favor the null model without this interaction. Finally, none of the other interactions were significant (all *p* ≥ .187).

#### Accuracy Analysis

Mean accuracy for each condition is displayed in **Figure S2** and tabulated in **Table S2**. Accuracy was subject to ceiling effects, such that the distribution of participants’ scores violated assumptions of normality (Shapiro-Wilk W = 0.82, *p* < .001) and homogeneity (Levene’s Test F = 26.3, *p* < .001) of variance. Results should therefore be interpreted with caution. There was a significant main effect of task sequence, with participants responding less accurately on switch (M_switch_ = 85.1%, 95% CI [83.1, 87.0]) than repeat trial (M_repeat_ = 91.1, 95% CI [89.9, 92.3]; F(1, 59) = 50.45, *p* < .001, *η_p_*^2^ = .461), and a significant main effect of stimulus congruency, with participants responding less accurately on incongruent (M_incongruent_ = 81.6%, 95% CI [79.4, 83.9]) compared to congruent trials (M_congruent_ = 94.5%, 95% CI [93.5, 95.5], F(1,59) = 145.92, *p* < .001, *η_p_*^2^ = .712). The interaction between task sequence and congruency was also significant (F(1,59) = 6.91, *p* = .011, *η_p_*^2^ = .103), as congruency effects were larger on switch (M_congruencyeffect_ = 14.6%, 95% CI [12.1, 17.1]) compared to repeat trials (M_congruencyeffect_ = 11.2%, 95% CI [8.7, 13.6]).

Recapitulating the RT analysis, there were significant interactions between block-wise switch proportion and task sequence (F(1,59) = 28.76, *p* < .001, *η_p_*^2^ = .328) and between block-wise congruency proportion and stimulus congruency (F(1,59) = 68.9, *p* < .001, *η_p_*^2^ = .539). Accuracy switch costs were smaller for the 75% switch condition (M_switchcost_ = 2.7%, 95% CI [0.6, 4.8]) compared to the 25% switch condition (M_switchcost_ = 9.3%, 95% CI [7.2, 11.4]), and the congruency effect was smaller for the 75% incongruent condition (M_congruencyeffect_ = 8.5%, 95% CI [6.2, 10.9]) compared to the 25% incongruency condition (M_congruencyeffect_ = 17.2%, 95% CI [14.8, 19.6]). Unlike the RT analysis, there was a significant interaction between the block-wise congruency proportion and task sequence, with larger switch costs for the 75% incongruent condition (M_switchcost_ = 7.2%, 95% CI [5.2, 9.2]) compared to the 25% incongruency condition (M_switchcost_ = 4.8%, 95% CI [2.8, 6.8]; F(1,59) = 4.51, *p* = .038, *η_p_*^2^ = .071). A Bayesian rmANOVA produced a best-fitting model of *task_sequence + congruency + LWPC + LWPS + task_sequence * congruency + task_sequence * LWPS + congruency * LWPC*. Compared to this best-fitting model, the model with the significant task sequence * LWPC had a BF_01_ = 2, suggesting very weak evidence in favor of the model without this interaction term. The interaction between stimulus congruency and switch proportion was not significant (F(1,59) = 0.88, *p* = .352, *η_p_*^2^ = .015), with the best model with this term having a BF_01_ = 6.7, suggesting moderate evidence against this interaction.

Finally, a significant three-way interaction was found between task sequence, LWPS, and LWPC (F(1,59) = 4.58, *p* = .037, *η_p_*^2^ = .072), due to a smaller effect of switch proportion on switch costs in the 25% incongruency condition (M = 4.8%, 95% CI [1.8, 7.8]) than in the 75% incongruency condition (M = 8.5%, 95% CI [5.5, 11.4]). However, this interaction term was not in any of the best fitting Bayesian rmANOVA models, and the best model with this interaction had a BF_01_ = 51.1, providing extremely strong evidence against it.

### Discussion

In Experiment 2, we sought to investigate if the same independence of adaptation in stability and flexibility we observed in the context of cued task switching in Experiment 1 also existed in the context of attentional set shifting, where participants have to move between attending to different features of a compound stimulus to respond correctly to a constant task rule. Crucially, we replicated Experiment 1’s key response time results: we observed robust effects of set shifting and congruency, as well as significant adaptation of these effects to changes in shift and congruency proportion over blocks of trials; however, we observed no effects of stability regulation on shift costs nor of flexibility regulation on congruency effects. Note that there were again numerical differences in switch costs between congruency proportions and in congruency effects between switch proportions in the direction predicted by a stability-flexibility tradeoff, but these differences were not reliable as indicated by the conventional ANOVA, and the corresponding interaction terms were again refuted by Bayesian analyses. Thus, just as in the domain of task switching, the regulation of cognitive stability and flexibility appear to be largely independent of each other in the domain of set shifting, too. Additionally, there was a significant three-way interaction, but this effect went in an opposite direction of the tradeoff perspective, and the Bayesian analysis strongly rejected these interaction terms, and they should therefore be interpreted very cautiously. While recapitulating the key RT results, the accuracy analysis also included a significant interaction between task sequence and congruency proportion indicative of a potential stability-flexibility tradeoff, but the Bayesian rmANOVA was unable to reliable distinguish between the best fitting with model without this term and the one with it, and the accuracy data violated distributional assumptions of the ANOVA. Taken together, we observed little evidence to support an obligatory yoking between stability and flexibility.

## Experiment 3

Experiment 1 and 2 examined cognitive stability and flexibility in the domains of task switching and set shifting, respectively, and both produced robust evidence in favor of independent mechanisms for regulating stability and flexibility. In Experiment 3, we attempted another conceptual replication of these findings in a protocol that modified the relationships between target and distracter information and responses from the first two experiments. In particular, one commonality between the protocols used in Experiments 1 and 2 is that they employed single, bivalent or compound stimuli and overlapping response sets. This aspect of these task designs maximizes the potential for interference between task rules (Experiment 1) and stimulus features (Experiment 2), as the “distracter” feature is both an integral part of the target stimulus and also shares responses with the target. In real life, however, we are often distracted by things that are extraneous to our current focus of attention and do not compete for identical responses (e.g., we might be distracted by our child calling us while trying to write a paper). In order to approximate this, perhaps more common scenario, in Experiment 3 we designed a hybrid task that crossed a task switching and a flanker task protocol (**Figure 5**). Here, the two tasks had different response sets, and distracters were not integral to the target stimuli but rather were spatially adjacent to those stimuli. As in the previous two experiments, flexibility was assessed via switch costs, and stability via flanker congruency effects, and their adaptation was gauged via manipulating switch and congruency proportions over blocks of trials.

**Figure 5:**
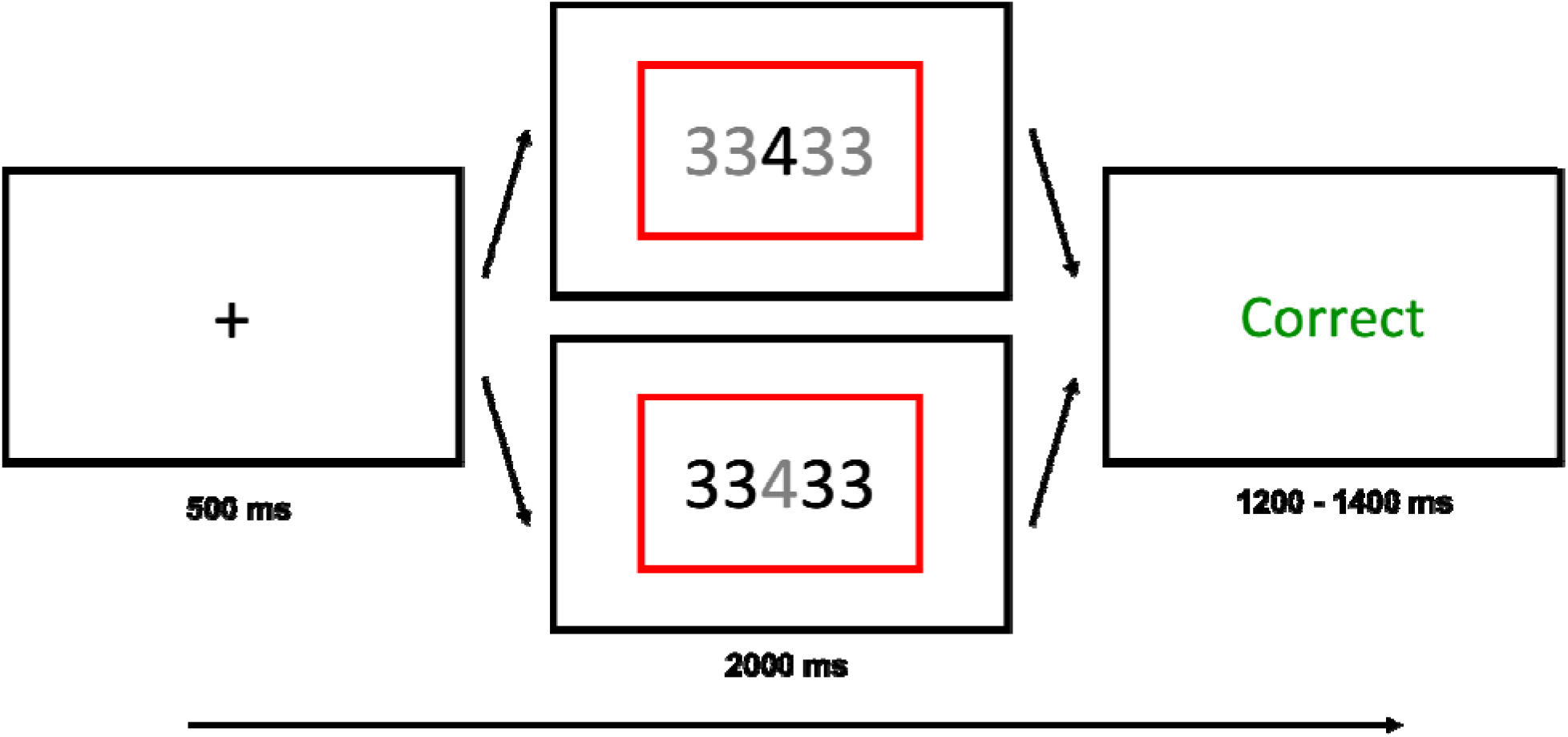
The trial structure for Experiment 3 consisted of an intertrial fixation cross preceding stimulus presentation with a colored rectangle (blue or red) cueing participants to either the magnitude or parity task, followed by accuracy feedback. The color of the numbers indicated which number participants needed to attend to (black = task relevant, grey = task irrelevant).

### Methods

#### Participants

63 participants completed the experiment on Amazon Mechanical Turk for financial compensation. 6 participants (9.5%) were excluded for failing to meet the accuracy criterion (>75% correct overall and 60% on catch trials), leaving a final sample of 57 participants (34 male, 23 female, 1 did not specify; age range: 22 - 58, mean: 36.63, SD: 8.36). The study was approved by the Duke University Campus Institutional Review Board.

#### Stimuli and Procedure

Task stimuli for Experiment 3 consisted of a single center digit (1 – 9 excluding 5), flanked on both sides by distractor digits (**Figure 5**). Participants alternated between categorizing the center digit’s parity (odd or even) or magnitude (greater or less than 5), based on the color of the rectangular frame. In order to increase the potential for interference from the distractor digits, on rare “catch trials” (12.5%) participants were cued to indicate the parity or magnitude of the distractors digits instead of the center digit (Armbruster-Genç et al., 2016). Because this renders the flankers task-relevant on some trials, the inclusion of these catch trials should generally increase participants’ attention to the distracter digits, and thus result in larger flanker interference (congruency) effects. Target identity (i.e., whether responding to center vs. distractor digits) was indicated based on the color of the digits, with black indicating the digit participants had to respond to, and the non-target digits shown in grey. Thus, on 87.5% of trials, the center digit was shown in black and the distracters in grey, and in 12.5% of trials (catch trials) it was the other way around.

Each trial began with a fixation cross (500 ms), followed by stimulus presentation (2000 ms or until response). Participants were required to indicate the target digit’s parity or its magnitude based on the color of a rectangular frame (blue or red) that surrounded the number. Color-to-task mapping was randomized across participants. Unlike in Experiment 1, the protocol used in Experiment 3 had non-overlapping response sets. Participants used their left hand index and middle finger on the ‘Z’ and ‘X’ keys, respectively, and their right hand index and middle finger on the ‘N’ and ‘M’ keys, respectively, to respond to the parity and magnitude task, respectively. Task-hand mapping and category-response mapping was randomized across participants. Trials were classified as congruent if the target digit and distractor digits were the same (e.g., a ‘4’ flanked by more ‘4’s) and as incongruent if they were different. The congruency effect again served as a measure of cognitive stability, with smaller congruency effects indicating a more stable task focus. To maximize this effect, distractor digits were chosen so as to be from the opposite category to the center digit in the cued task dimension. For example, if the center digit was a ‘4’ on an incongruent parity trial, the distractor digits would be either 1 or 3, as these digits do not match the center digit in the parity dimension. Trials were also classified as repeat or switch trials based on whether or not the task cue changed between trials (i.e., a parity trial following a magnitude trial would be a switch trial). The switch cost again served as a measure of cognitive flexibility, with smaller switch costs indicating a more flexible state.

The practice phase consisted of 4 blocks of 16 trials each. The first practice block consisted entirely of parity trials and the second of magnitude trials (practice order randomized). The third practice block consisted of a mix of parity and magnitude tasks. The fourth and final block introduced catch trials, where participants had to respond to the distractors instead of the target on 12.5% of trials. The main experiment consisted of 4 blocks of 128 trials each, resulting in an equivalent trial count to Experiments 1 and 2. We again varied the relative frequency of incongruent stimuli (25% vs 75%) and the frequency of switch trials (25% vs 75%) across blocks, with block order counterbalanced across participants using a Latin square design. Thus, Experiment 3 conformed to the same factorial design as Experiments 1 and 2, and the analysis strategy was also identical. We expected to reproduce standard congruency, proportion congruent, task switch, and proportion switch effects. The main question of interest was again whether congruency effects would interact with the proportion switch factor and whether switch costs would interact with the proportion congruent factor.

### Results

#### Trial exclusions

Trial exclusions for Experiment 3 were the same as for Experiments 1 and 2, including removal of incorrect trials (7.0%), 3 times standard deviation RT trials (0.7%), and trials faster than 300 ms or slower than 2000 ms (<0.1%). We also removed catch trials from the experiment (12.5%).

#### RT Analysis

The mean RT for each cell of the experimental design is displayed in **Figure 6a**, and descriptive statistics can be found in **Table S1**. There was a significant main effect of task sequence, reflected in slower RTs for switch trials (M_switch_ = 1027 ms, 95% CI [994, 1061]) compared to repeat trials (M_repeat_ = 902 ms, 95% CI [868, 936]; F(1, 56) = 209.62*, p* < .001, *η_p_*^2^ = .789), and a significant main effect of stimulus congruency, due to slower RTs for incongruent (M_incongruent_ = 983ms, 95% CI [950,1016]) compared to congruent stimuli (M_congruent_ = 946ms, 95% CI [913, 979]; F(1, 56) = 73.33, *p* < .001, *η_p_*^2^ = .567). There was also a significant interaction between task sequence and stimulus congruency (F(1,56) = 9.10, *p* = .004, *η_p_*^2^ = .140), as congruency effects were smaller on switch (M_congruencyeffect_ = 36 ms, 95% CI [21, 51]) compared to repeat trials (M_congruencyeffect_ = 57 ms, 95% CI [42, 72]). Note that this finding is opposite the effect sometimes seen in the literature (e.g., Kiesel et al., 2010; Wendt & Kiesel, 2008). Finally, we also observed a main effect of LWPS (F(1,56) = 44.34, *p* < .001, *η_p_*^2^ = .442) due to slower RTs in the 75% switch condition (M_75_ = 984, 95% CI [951, 1017]) than the 25% switch condition (M_25_ = 945, 95% CI [912, 978]), but the main effect of LWPC was not significant (F(1,56) = 0.50, *p* = .483, *η_p_*^2^ = .009).

**Figure 6:**
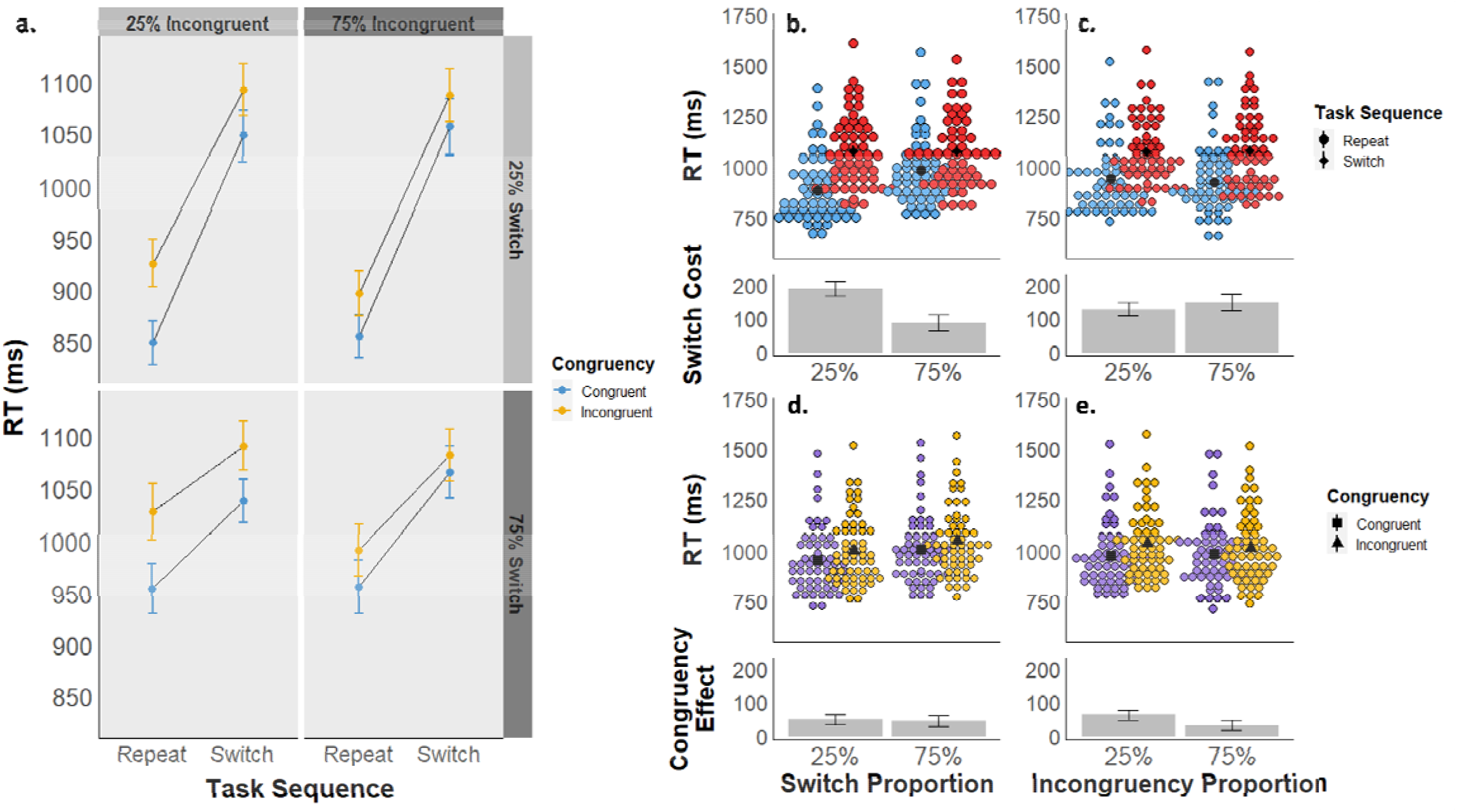
**a)** Experiment 3 mean reaction times are displayed as a function of task sequence (switch vs. repeat), stimulus congruency (congruent vs. incongruent), switch proportion (25% vs. 75%), and incongruency proportion (25% vs. 75%). **b-e)** Experiment 3 mean RTs are displayed as a function of task sequence (panels b and c) and congruency (d and e), collapsed across block-wise switch proportions (b and d) and block-wise congruency proportions (c and e). The upper graphs in each panel depict mean RTs by subject for each condition, and the lower graphs depict the mean RT difference between conditions.

There was a significant interaction between task sequence and the block-wise switch proportion (i.e., 25% vs. 75% switch trials; F(1,56) = 128.97, *p* < .001, *η_p_*^2^ = .697). Switch costs were smaller in the 75% switch proportion condition (M_switchcost_ = 87 ms, 95% CI [65, 109]) than in the 25% switch proportion condition (M_switchcost_ = 189 ms, 95% CI [167, 211]; **Figure 6b**). We also observed a significant interaction between congruency and the block-wise congruency proportion (F(1,56) = 11.98, *p* < .001, *η_p_*^2^ = .176). Congruency effects were smaller in the 75% incongruency proportion condition (M_congruencyeffect_ = 31 ms, 95% CI [16, 46]) than in the 25% incongruency proportion condition (M_congruencyeffect_ = 62 ms, 95% CI [47, 77]; **Figure 6e**).

Our primary effect of interest was again the interaction between block-wise switch proportion and congruency effects, and conversely, between block-wise congruency proportion and switch costs. Unlike Experiment’s 1 and 2, there was a significant interaction between the congruency proportion and task sequence (F(1,56) = 5.40, *p* = .024, *η_p_*^2^ = .088), due to larger switch costs in the 75% incongruency condition (M_switchcost_ = 148 ms, 95% CI [125, 171]) than in the 25% incongruency condition (M_switchcost_ = 128 ms, 95% CI [106, 151]; **Figure 6c**). However, a bayesian rmANOVA found a best fitting model of *task_sequence + congruency + LWPS + task_sequence * congruency + task_sequence * LWPS.* Compared to this model, the best fitting model with the significant task sequence * LWPC interaction term had a BF_01_ = 12.8, suggesting strong evidence against this term. Note that the best-fitting model did not have the expected congruency * LWPC interaction, and the model that did had a BF_01_ = 4.3 compared to it, despite this interaction also being significant. The interaction between switch proportion and stimulus congruency was not significant (F(1,56) = 0.03, *p* = .870, *η_p_*^2^ < .001; **Figure 6d**), and the best fitting Bayesian rmANOVA model with this term had a BF_01_ = 9.6. Finally, none of the higher order interactions were significant (all *p* ≥ .286).

#### Accuracy Analysis

Mean accuracy for each condition is displayed in **Figure S3** and tabulated in **Table S2**. Accuracy was subject to ceiling effects, such that the distribution of participants’ scores violated assumptions of normality (Shapiro-Wilk W = 0.8, *p* < .001) and homogeneity (Levene’s Test F = 10.4, *p* < .001) of variance. Results should therefore be interpreted with caution. There was a significant main effect of task sequence, with participants responding more accurately on repeat (M_repeat_ = 94.3%, 95% CI [92.7, 95.9]) compared to switch trials (M_switch_ = 91.2%, 95% CI [89.6, 92.7]; F(1, 56) = 22.41, *p* < .001, *η_p_*^2^ = .286), but the main effect of stimulus congruency was not significant, with similar accuracies for congruent (M_congruent_ = 92.9%, 95% CI [91.4, 94.4]) and incongruent trials (M_incongruent_ = 92.6%, 95% CI [91.1, 94.1]; F(1,56) = 0.40, *p* = .528, *η_p_*^2^ = .007). There was again a significant main effect of the block-wise switch proportion (F(1,56) = 5.08, *p* = .028, *η_p_*^2^ = .083), due to higher accuracies in the 25% switch condition (M_25_ = 93.3%, 95% CI [92.0, 94.7]) than the 75% switch condition (M_75_ = 92.1%, 95% CI [90.5, 93.8]), but no main effect of the block-wise congruency proportion (F(1,56) = 0.96, *p* = .331, *η_p_*^2^ = .017).

There was no significant interaction between the block-wise switch proportion and task sequence (F(1,56) = 0.83, *p* = .366, *η_p_*^2^ = .015), nor the block-wise congruency proportion and stimulus congruency (F(1,56) = 0.90, *p* = .347, *η_p_*^2^ = .016) in performance accuracy. There was a significant interaction between the block-wise switch proportion and stimulus congruency (F(1,56) = 4.29, *p* = .043, *η_p_*^2^ = .071), due to a larger congruency effect in the 25% switch condition (M_25_ = 0.7%, 95% CI [-0.6, 2.1]) than the 75% switch condition (M_75_ = -1.3%, 95% CI [-2.7, 0.1]). However, this interaction term was not part of the best-fit model of the Bayesian rmANOVA, and the best model including a significant *congruency * LWPS* interaction had a BF_01_ = 15.8, providing very strong evidence against this term. The interaction between the block-wise congruency proportion and task sequence was not significant (F(1,56) = 0.26, *p* = .611, *η_p_*^2^ = .005). Importantly, the best-fitting model produced by the Bayesian rmANOVA included only the *task_sequence* term, suggesting that variance in performance accuracy in this experiment was primarily driven by switch costs.

### Discussion

Experiment 3 again assessed the inter-dependence of control over cognitive stability and flexibility, but the design physically separated the target and distracter stimuli and the response sets between tasks. This resulted in smaller congruency effects, even with additional catch trials designed to maximize them. We again replicated the expected LWPS and LWPC effects in response times, though evidence for the latter was scant in the Bayesian model of the data. We additionally observed a significant interaction between task sequence and congruency proportion, unlike in Experiments 1 and 2. Although this interaction suggests some degree of interdependence between cognitive stability and flexibility, the Bayesian rmANOVA provided strong evidence against the inclusion of this effect in the best-fitting model of the RT data, and the same was true for the accuracy results). Overall, the results of Experiment 3 were less conclusive than those of Experiments 1 and 2, primarily due to less robust effects of conflict and conflict-control, but again we found no convincing evidence in favor of an obligatory inverse yoking between stability and flexibility.

## General Discussion

This study sought to rigorously test the widely held assumption that cognitive stability and flexibility are reciprocal. To this end, we acquired simultaneous and independent measurements of stability and flexibility combined with block-wise proportion manipulations known to induce strategic adjustments in stability and flexibility (i.e., the LWPC and LWPS effects, respectively). If stability and flexibility were opposing poles on a stability-flexibility continuum, then dynamic adjustments in favor of greater flexibility, as indexed by decreasing switch costs, should necessarily result in commensurate decreases in cognitive stability, as indexed by increasing cross-task congruency effects, and vice versa. Across three experiments, we found robust evidence against the stability – flexibility tradeoff assumption. We reproduced the expected adjustments in flexibility and stability caused by trial proportion manipulations of switching and congruency, but critically, these adjustments had little influence on the converse indices of stability and flexibility, respectively. Experiment 1 examined the interdependency assumption of the stability-flexibility tradeoff using a classic task switching protocol, with cognitive flexibility operationalized as switching between task rules and stability as preventing cross-task interference caused by overlapping response sets. We found robust evidence that increases in task focus do not influence task-switching ability and that readiness to switch tasks does not influence cross-task interference. Experiment 2 extended these findings by investigating cognitive stability and flexibility in terms of attentional set shifting. Here again we found evidence for the independency of stability and flexibility. Finally, Experiment 3 employed a hybrid task combining task switching and flanker interference. Congruency effects were substantially smaller in this paradigm, rendering conclusions less robust, but ultimately we again found little convincing evidence that increased flexibility would necessarily reduce stability reduce stability, and vice versa.

Support against the reciprocity view came from our repeated measures ANOVA results as well as the Bayesian counterpart, which allowed us to quantify evidence in favor of null hypotheses, as well as to evaluate relative fit of models with or without the theoretically important interaction terms. In Experiments 1 and 2, we found significant interactions between task sequence and switch proportion (i.e., LWPS effect) and congruency and congruency proportion (i.e., LWPC effect). By contrast, even though the pattern of means went in the direction of a tradeoff, the interactions between task sequence and congruency proportion and between congruency and switch proportion were not significant, and Bayesian model comparisons provided robust evidence against these interactions. Similarly, while some of the accuracy rmANOVAs and the RT data in Experiment 3 suggested a degree of reciprocity between stability and flexibility, moderate to strong Bayes Factor evidence spoke against the inclusion of these interaction terms in the best-fit models of the data. Taken together, the present results therefore converged on a view counter to the idea that stability and flexibility are subject to an obligatory or structural tradeoff, although – as we discuss below – this does not rule out the possibility of a functional tradeoff under many circumstances.

Together, these results provide a proof-of-principle demonstration that stability and flexibility can be regulated independently from one another. In turn, this refutes the possibility of a structural (non-malleable) reciprocity between the cognitive processes supporting stability and flexibility, as implied by the idea of a one-dimensional stability-flexibility continuum or a single meta-control parameter governing a stability/flexibility tradeoff. Instead, the current results suggest that stability- and flexibility-supporting processes should be conceptualized as lying on separate, orthogonal axes, thus making it possible to simultaneously engage a strong focus on the current task set while also being ready to swiftly switch sets (as was the case in the high LWPC/high LWPS conditions in the current experiments).

What form may those putatively independent mechanisms take? A substantial literature supports the idea that on-task focus (stability) is regulated via a control feedback loop that tracks ongoing variations in task difficulty (assessed via the proxy of conflict in information processing) and implements a commensurate up- or down-regulation of the strength of task set representations in working memory (Botvinick et al., 2001; for recent reviews, see Chiu & Egner, 2019; Egner, 2017). Thus, encountering a sequence of mostly congruent trials would result in a relatively low level of conflict-driven task focus, whereas a sequence of mostly incongruent trials would promote a strongly activated task set, leading to the LWPC effects (e.g., Bugg & Chanani, 2011; Spinelli & Lupker, 2020). As noted in the Introduction, a commonly proposed mechanism of task-set updating is a gating process whereby the gate to working memory (holding the task set) is opened in response to cues of changing incentives or task requirements, in order to switch out task sets (Frank et al., 2001; Hazy et al., 2006). Thus, one plausible way in which switch-readiness in the present experiments could be adjusted would be by some form of priming of this gate (corresponding to a lowering of the updating threshold, cf. Goschke, 2003; 2013), such that it can be opened more easily in scenarios where many recent trials entailed task switches. If this were the mechanism for adapting flexibility in the present experiments, our results would suggest that the ease of gate-opening, and thus of switching task set, is independent from the (conflict-driven) strength with which the relevant task set will be enforced. This seems plausible given the notion that task set activation and updating may be reliant on different neuroanatomical loci, the prefrontal cortex and basal ganglia, respectively (Collins et al., 2000; Crofts et al., 2001; Durstewitz et al., 2000; Roberts et al., 1994).

An alternative possibility is that, rather than affecting the updating threshold, high switch readiness between two tasks could be achieved by keeping both task sets in working memory simultaneously (Dreisbach & Fröber, 2019; Fröber et al., 2018). This proposal can account for a lack of transfer of switch proportion effects to probabilistically unbiased (Siqi-Liu & Egner, 2020) or novel tasks (Sabah et al., 2019), and has been explicitly proposed to mediate the type of frequency-driven adjustments in switch costs observed in the present experiments (Dreisbach & Fröber, 2019; Fröber et al., 2018). Under this assumption, the currently relevant task set would of course still have to be subject to some form of selection within working memory in order to be applied when cued, likely by being in the focus of internal attention (see Cowan, 2017; Oberauer & Hein, 2012). If this were the mechanism for adapting flexibility in the present experiments, our results would suggest that the efficiency of shifting the focus of attention between two task sets in working memory is independent from the strength with which the selected task set is being enforced based on the recent history of cross-task interference. This conjecture concerning internal selection in working memory could make for an interesting target of future studies.

We note, however, that the fact that cognitive stability and flexibility are not inversely yoked in an obligatory fashion - as demonstrated in the present study - does not rule out the possibility of a temporary, functional yoking of processes supporting stability and flexibility under circumstances where the task environment preferentially incentivizes either stability or flexibility. In particular, if one assumes that people are generally loath to invest cognitive effort if they can avoid it, as supported by much recent evidence (Botvinick & Braver, 2015; Kool & Botvinick, 2018; Shenhav et al., 2013; Westbrook & Braver, 2015), then a scenario where either only stability or only flexibility is incentivized is liable to produce the appearance of reciprocity, as we discuss next.

As reviewed in the introduction, most prior work in this literature has been consistent with a stability-flexibility tradeoff (Braem, 2017; Dreisbach & Wenke, 2011; Hefer & Dreisbach, 2017; Meiran & Kessler, 2008; Muhmenthaler & Meier, 2021; Müller et al., 2007; Musslick et al., 2018) without directly testing for it. We propose that one key reason for the appearance of a tradeoff might be that prior studies have focused primarily on task manipulations that would promote either stability or flexibility, but not both. Since cognitive control is effortful (Shenhav et al., 2013, 2016) and only implemented on an as-needed basis (Bugg, 2014), context manipulations that, for example, incentivize greater flexibility might result in a decrease in stability not because of an inherent, mechanistic stability-flexibility tradeoff but rather because simultaneously maintaining a stable task focus would require even greater cognitive control engagement, and that effort is not incentivized by the task. For instance, as described previously, Braem (2017) found that selectively rewarding task switches during cued task switching increased subsequent congruency effects. This result is consistent with the idea of a stability-flexibility tradeoff; however, that study could not ascertain whether the observed stability decrease (increased congruency effects) resulted from an obligatory tradeoff or instead simply reflected a relaxation of cognitive stability due to stability not being incentivized. Additionally, a strategic reduction of effort investment in cognitive stability may have been encouraged by the very fact that there was a high value placed on engaging cognitive control for flexibility. In other words, in light of the current results, we suggest that the findings by Braem (2017) and similar studies (Dreisbach & Wenke, 2011) may reflect a preferential effort-investment in flexibility-supporting processes and an accompanying effort-divestment from stability-supporting processes due to *expected value-of-control* computations (c.f., Shenhav et al., 2013) rather than due to a structural reciprocity between these processes. The reverse scenario would of course apply to conditions that selectively incentivize cognitive stability – here, the effort investment in task focus would be accompanied by a relaxation of flexibility since the latter is not promoted by the task structure.

In line with this argument, one major reason why previous studies have not detected independent regulation of stability and flexibility likely lies with the fact that they have studied these constructs in a static fashion, using a task environment with fixed incentives. The prime example is the assessment of the interaction between mean switch costs and congruency effects in experiments with a constant switch and congruency rate (typically both held at 50%; (Wendt & Kiesel, 2008). The commonly observed finding of increased cross-task congruency effects on switch compared to repeat trials here merely reflects a snapshot of the relationship between these measures given one particular, fixed level of flexibility and stability. Only by dynamically and independently varying demands on, or incentives for, stability and flexibility is it possible to properly gauge their degree of mutual dependence. When doing so in the present experiments, we clearly document that in conditions where stability and flexibility are both called for, people are in fact able to increase task shielding and switch-readiness at the same time. Future studies are required to further corroborate these results, ideally by taking other approaches for independently manipulating stability and flexibility demands. For example, according to the orthogonal axes view of stability and flexibility espoused here, we would expect to see equivalent results to the current ones under independent manipulations of reward for task switching and shielding operations. Another fruitful line of research would be to adopt the current protocol to neuroimaging; on the basis of the current behavioral results, we would expect to observe distinct neural mechanisms to mediate adaptation to varying demands on stability and flexibility.

Another possible reason that switch costs and cross-task congruency effects have often been found to be inversely related under conditions of stable incentives (i.e., a 50% switch and congruency rates throughout an experiment), but not when examining strategic adjustments in stability and flexibility (in the present study), might be that these conditions encourage distinct reactive versus proactive modes of applying cognitive control. According to the dual mechanisms of control framework (Braver, 2012), cognitive control can be engaged in two distinct ways, one in which goal-relevant information is maintained in (contextually cued) anticipation of cognitive demands (proactive control), and one in which control is recruited on an as-needed basis in direct response to encountering demands (reactive control). A set-up where neither distracter interference nor task switching are incentivized or anticipated (i.e., 50% switch and congruency rates) may induce a relatively reactive mode of control, and this in turn could lead to temporary resource competition or processing bottlenecks when task switching and shielding operations have to be engaged simultaneously, especially when task cues and target stimuli are presented at the same time or with a very short cue-to-target interval. By contrast, a situation where the task statistics allow for a more proactive strategy due to a low or high likelihood of switching or incongruent distracters, the underlying control processes could be engaged in a fashion that reduces temporal overlap and competition between them. Some evidence in support of this idea comes from task switching experiments showing that a longer cue-to-target interval reliably reduces switch costs (Hoffmann et al., 2003; Kiesel et al., 2010; Kiesel & Hoffmann, 2004; Koch, 2001; Meiran et al., 2000; Monsell et al., 2003) but does not result in a commensurate increase in congruency effects (Allport et al., 1994; Meiran, 1996; Rogers & Monsell, 1995). Thus, being able to engage in a more proactive task-set updating strategy appears to uncouple the functional yoking of switch costs and congruency effects that is typically observed at short or zero cue-to-target intervals. Note that this finding does of course also speak against the assumption of a single mechanism or control parameter determining both stability and flexibility.

Finally, it is worth noting that the observed flexibility adaptation in all three experiments (i.e., reduced switch costs in the 75% switch proportion compared to the 25% switch proportion; the LWPS effect) is driven primarily by slower task repetition trials in the 75% switch condition, rather than by faster task switch trials (Figures 1, 2, and 3). This data pattern is quite common in LWPS studies (e.g., Dreisbach & Haider, 2006; Siqi-Liu & Egner, 2020) but is nevertheless counterintuitive, since one might expect faster switch trial performance when switching more frequently. One plausible explanation for this, put forward by Bonnin and colleagues (2011), is that a potential speed-up on switch trials in high proportion switch blocks may be masked by backward inhibition effects (Mayr & Keele, 2000), the finding that people tend to be slower to switch back to a task that they only just switched away from (e.g., an ABA task sequence) than when switching to a less recently abandoned task (e.g., a CBA task sequence). Since the high proportion switch blocks in the current experiments entail more frequent switches back to the previous task than the low proportion switch blocks, backward inhibition effects may overshadow any RT benefits of increased switch preparation on switch trial performance in the former.

In conclusion, using a novel task design to independently manipulate demands on task shielding and task switching, we here show, for the first time, that cognitive flexibility and stability are not reciprocal but rather rely on separable mechanisms that can be independently dialed up or down. Thus, the widely held assumption of an obligatory tradeoff between cognitive stability and flexibility appears to be unfounded. We propose that instances where a stability/flexibility tradeoff is observed do not reflect a structural inverse yoking of stability- and flexibility-supporting processes, but rather an independent recruitment of such processes on the basis of the expected value of their engagement.

