## Supplemental Tables and Figures for "No need to choose: independent regulation of cognitive stability and flexibility challenges the stability-flexibility tradeoff"

**Supplementary Materials**

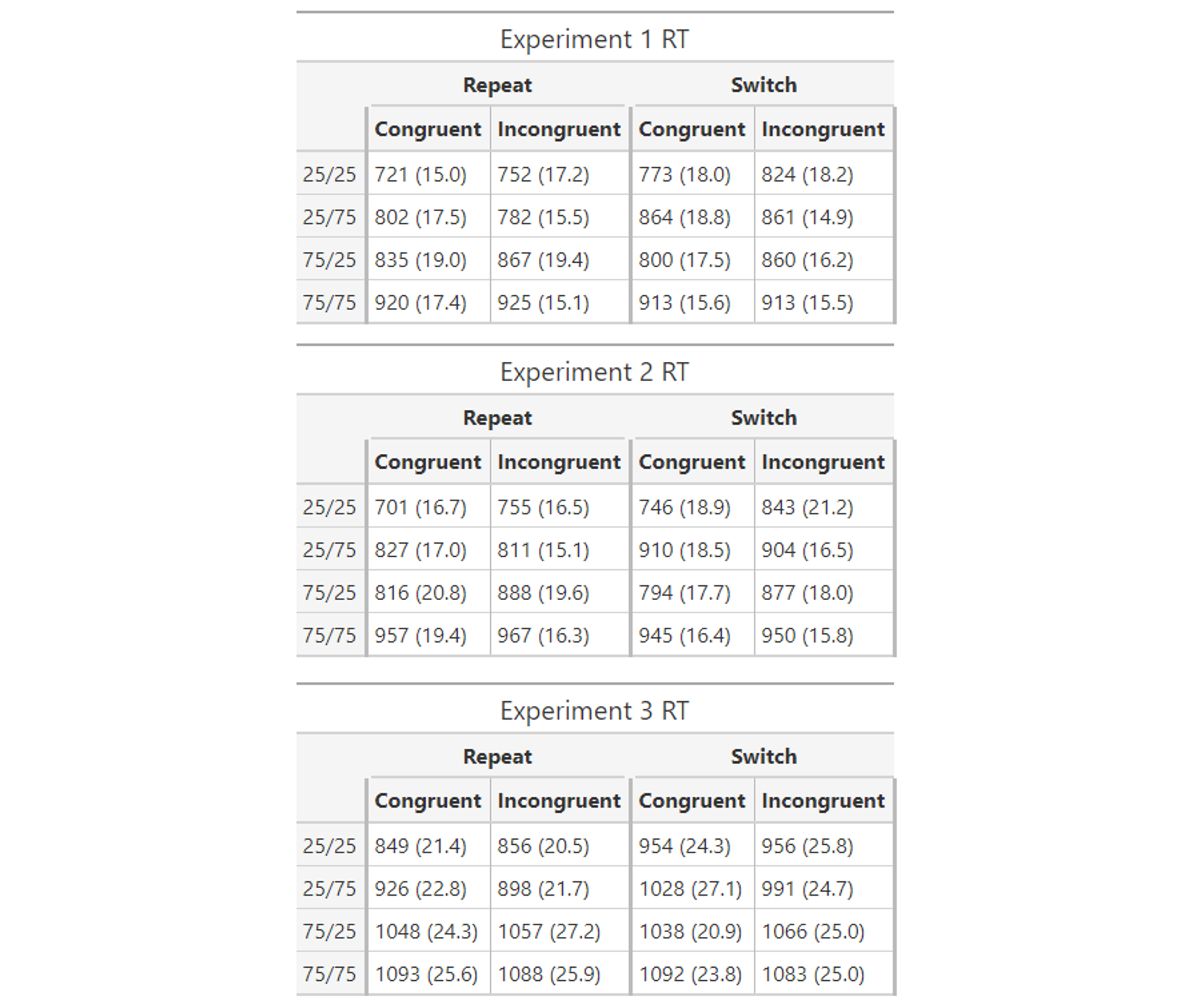

**Table S1:** Mean reaction times (SEM) by 2 x 2 x 2 x 2 factorial design. 25/25, 25/75, etc. refer to block-wise switch proportion / block-wise incongruency proportion.

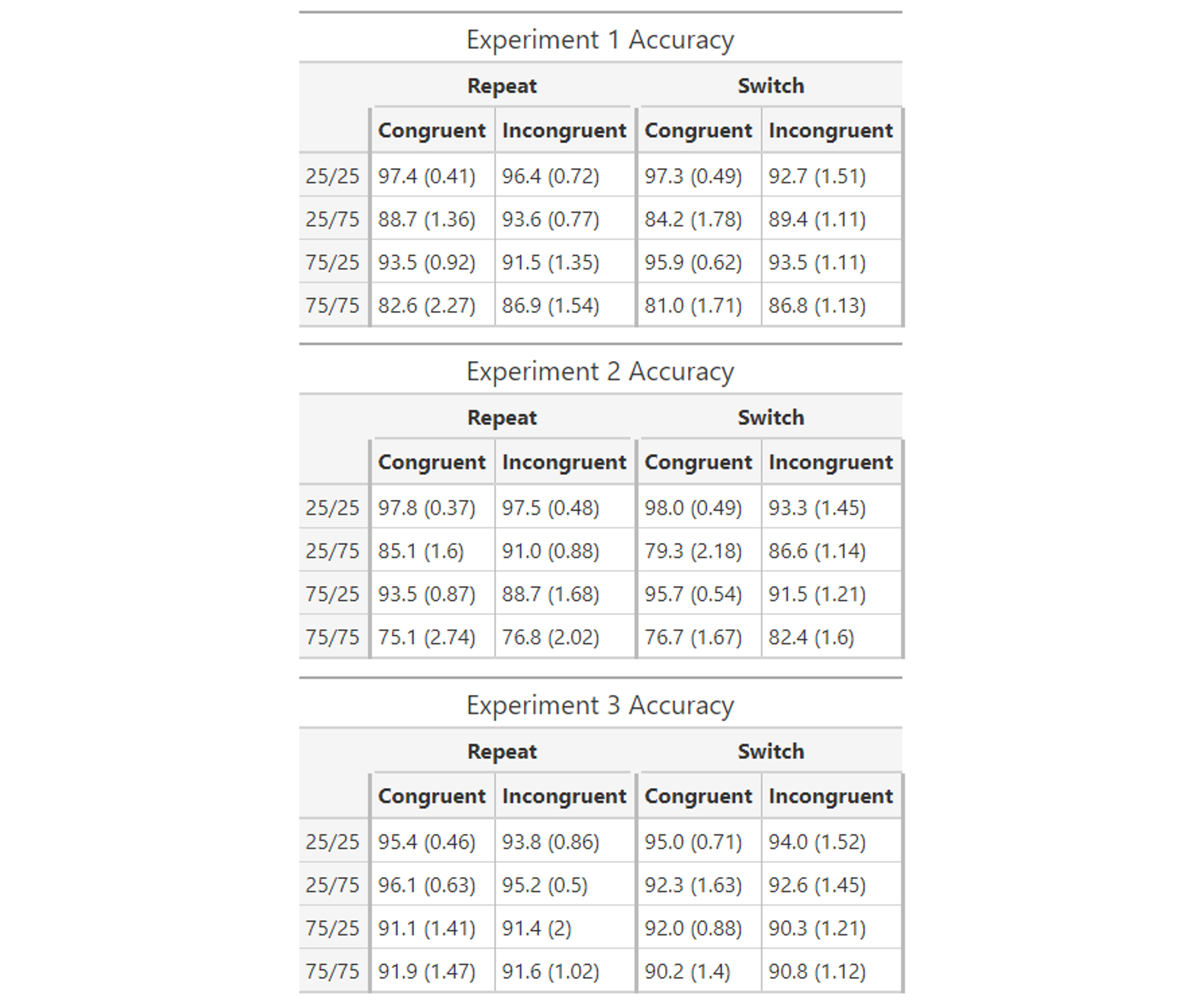

**Table S2:** Mean accuracies (SEM) by 2 x 2 x 2 x 2 factorial design. 25/25, 25/75, etc. refer to block-wise switch proportion / block-wise incongruency proportion.

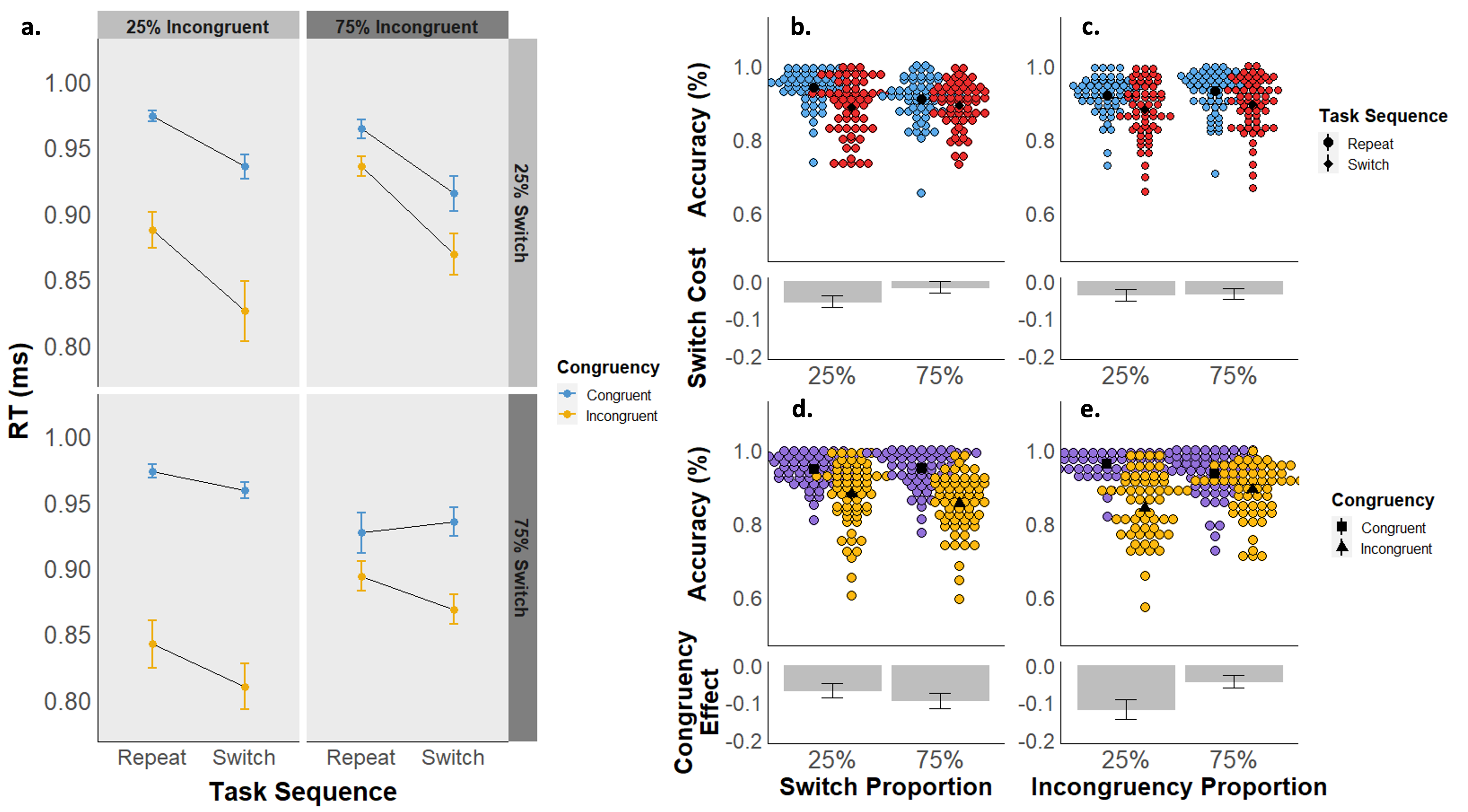

**Figure S1**: ***a)*** *Experiment 1 mean accuracies are displayed as a function of task sequence (switch vs. repeat), stimulus congruency (congruent vs. incongruent), switch proportion (25% vs. 75%), and incongruency proportion (25% vs. 75%).* ***b-e)*** *Experiment 1 mean accuracies are displayed as a function of task sequence (panels b and c) and congruency (d and e), collapsed across block-wise switch proportions (b and d) and block-wise congruency proportions (c and e). The upper graphs in each panel depict mean accuracy by subject for each condition, and the lower graphs depict the mean accuracy difference between conditions.*

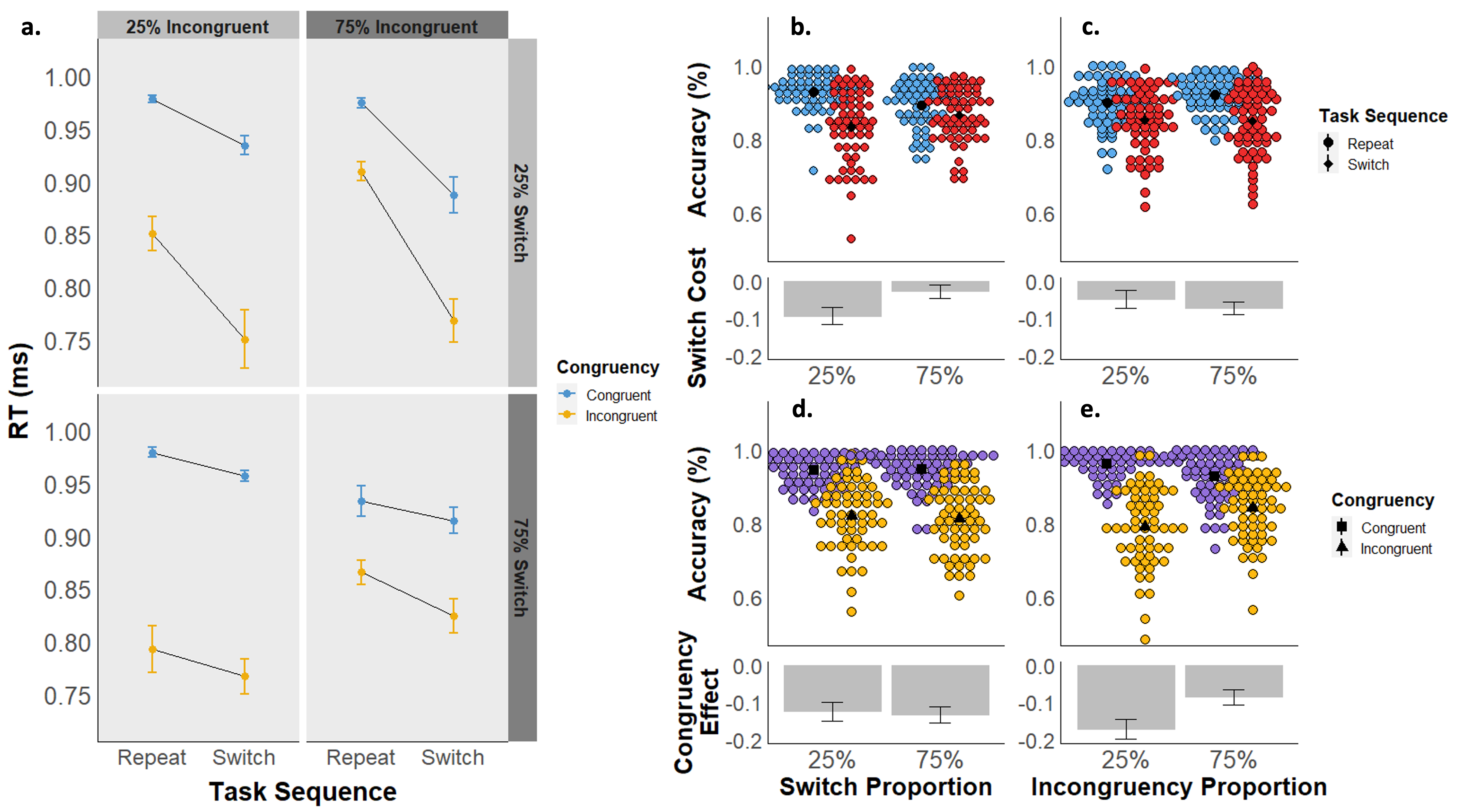

**Figure S2**: ***a)*** *Experiment 2 mean accuracies are displayed as a function of task sequence (switch vs. repeat), stimulus congruency (congruent vs. incongruent), switch proportion (25% vs. 75%), and incongruency proportion (25% vs. 75%).* ***b-e)*** *Experiment 2 mean accuracies are displayed as a function of task sequence (panels b and c) and congruency (d and e), collapsed across block-wise switch proportions (b and d) and block-wise congruency proportions (c and e). The upper graphs in each panel depict mean accuracy by subject for each condition, and the lower graphs depict the mean accuracy difference between conditions.*

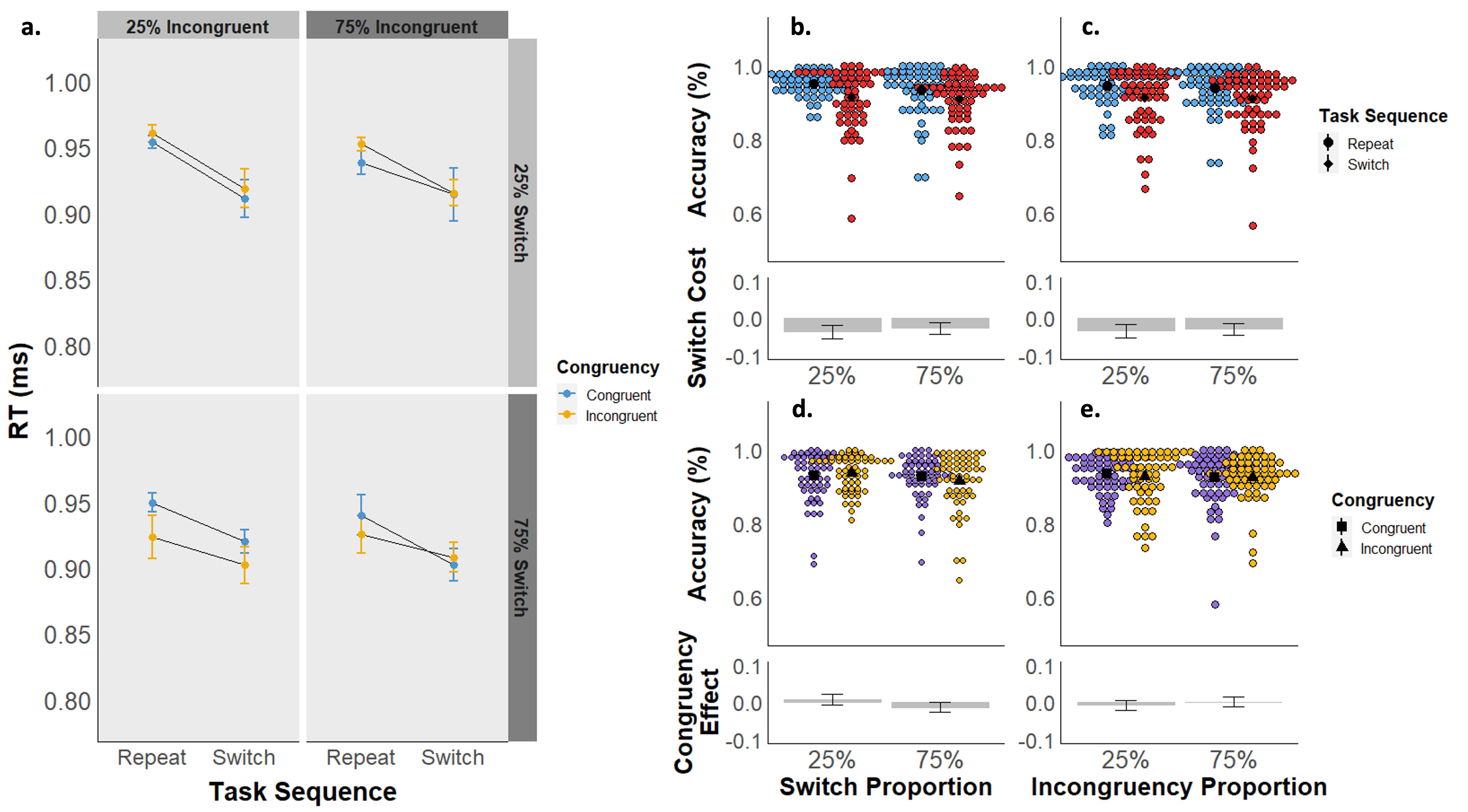

**Figure S3**: ***a)*** *Experiment 3 mean accuracies are displayed as a function of task sequence (switch vs. repeat), stimulus congruency (congruent vs. incongruent), switch proportion (25% vs. 75%), and incongruency proportion (25% vs. 75%).* ***b-e)*** *Experiment 3 mean accuracies are displayed as a function of task sequence (panels b and c) and congruency (d and e), collapsed across block-wise switch proportions (b and d) and block-wise congruency proportions (c and e). The upper graphs in each panel depict mean accuracy by subject for each condition, and the lower graphs depict the mean accuracy difference between conditions.*
